# Genomic sequencing confirms absence of introgression despite past hybridisation between a common and a critically endangered bird and its common congener

**DOI:** 10.1101/2020.09.28.316299

**Authors:** Natalie J. Forsdick, Denise Martini, Liz Brown, Hugh B. Cross, Richard F. Maloney, Tammy E. Steeves, Michael Knapp

## Abstract

Genetic swamping resulting from interspecific hybridisation can increase extinction risk for threatened species. The development of high-throughput and reduced-representation genomic sequencing and analyses to generate large numbers of high resolution genomic markers has the potential to reveal introgression previously undetected using small numbers of genetic markers. However, few studies to date have implemented genomic tools to assess the extent of interspecific hybridisation in threatened species. Here we investigate the utility of genome-wide single nucleotide polymorphisms (SNPs) to detect introgression resulting from past interspecific hybridisation in one of the world’s rarest birds. Anthropogenic impacts have resulted in hybridisation and subsequent backcrossing of the critically endangered Aotearoa New Zealand endemic kakī (black stilts; *Himantopus novaezelandiae*) with the non-threatened self-introduced congeneric poaka (Aotearoa New Zealand population of pied stilts, *Himantopus himantopus leucocephalus*), yet genetic analyses with a limited set of microsatellite markers revealed no evidence of introgression of poaka genetic material in kakī, excluding one individual. We use genomic data for ∼63% of the wild adult kakī population to reassess the extent of introgression resulting from hybridisation between kakī and poaka. Consistent with previous genetic analyses, we detected no introgression from poaka into kakī. These collective results indicate that, for kakī, existing microsatellite markers provide a robust, cost-effective approach to detect cryptic hybrids. Further, for well-differentiated species, the use of genomic markers may not be required to detect admixed individuals.

## 1 Introduction

Growing evidence from genetic and genomic data indicates that interspecific hybridisation (hereafter, hybridisation) has been integral in the evolutionary history of many species (Mallet, 2008), challenging existing perceptions of the intrinsic value of hybrids and hybrid species, and further highlighting the complexity of conservation policy relating to them (Haig & Allendorf, 2006; Jackiw et al., 2015; Wayne & Shaffer, 2016). Hybridisation may improve individual fitness, population resilience and adaptive potential (Arnold, 1997; Dowling & Secor, 1997; Mallet, 2007; Seehausen, 2004), and as such hybridisation between closely related species or subspecies has been proposed as a conservation management tool to assist genetically depauperate threatened species by introducing novel genetic variation to increase genetic diversity and improve fitness (Arnold, 2016; Harrisson et al., 2016; Ingvarsson, 2001; 65 Mallet, 2005). Nevertheless, potential tradeoffs resulting from conservation management of hybrids and hybridisation – especially recent human-induced hybridisation between threatened endemic and non-threatened non-endemic congeners – warrants careful ethical and practical consideration, balancing conservation priorities alongside the ecological, social, and economic costs-benefits of hybrids (e.g., Hamilton & Miller, 2016; Estévez et al., 2015; Schlaepfer et al., 2011).

Conservation is an inherently values-based discipline. From a conservation perspective, threatened endemic species are valued over non-threatened non-endemic species due to their rarity and their known (or perceived) ecological importance, which generally leads to ethical and moral obligations to conserve them, especially if they are also culturally significant species (Booth et al., 2011; Courchamp et al., 2006; Maguire & Justus, 2008; Richardson & Loomis, 2009). However, the potential conservation value of hybrids should not be viewed as static, especially as altered species ranges increase the prevalence of hybrids (Chunco, 2014), and the inclusion of genomic data continues to improve our understanding of the impacts of hybridisation (vonHoldt et al., 2018). Further, there are growing calls for researchers identify and avoid unscientific and biased perspectives on hybrids and hybridisation in conservation (Draper et al., 2021; Hirashiki et al., 2021).

Regardless of the perceived value of hybrids, hybridisation negatively impacts threatened species recovery through the misplaced reproductive efforts of interspecific breeding (Allendorf et al., 2001). This can reduce the reproductive outputs of species of conservation concern by demographic swamping (Allendorf et al., 2001; Wolf et al., 2001). Another potential impact of hybridisation is when subsequent backcrossing to the parental species incorporates genetic material from one species into the genome of another, known as introgression (Rhymer & Simberloff, 1996). Negative impacts of introgression may include outbreeding depression, where the breakdown of coadapted gene complexes or the introduction of maladaptive traits results in the decreased fitness of hybrid offspring (Arnold, 1997; Edmands, 2007; Lynch, 1991), and, at its most extreme, may result in extinction-by-hybridisation (Allendorf et al., 2001; Fitzpatrick et al., 2010; Quilodrán et al., 2018; Rhymer & Simberloff, 1996; Riley et al., 2003; Taylor et al., 2006; Todesco et al., 2016).

Genetic tools may be employed to assist conservation management programmes in assessing the extent and impacts of hybridisation, and for identification of cryptic hybrid offspring morphologically indistinguishable from parental types (Chan et al., 2006; Ma & Lambert, 1997; Milián-García et al., 2015; Pierpaoli et al., 2003). However, to date, most conservation-relevant studies have used a small number of genetic markers (e.g., microsatellites) that may not be representative of genome-wide diversity, particularly among threatened species where population bottlenecks have left populations genetically depauperate (Taylor, 2015; Taylor et al., 2015; Väli et al., 2008). Over the past decade, genomic sequencing technologies have progressed, and rapidly declining costs now enable the sequencing and assembly of complete genomes for threatened non-model organisms (e.g., Li et al., 2010; Sutton et al., 2018), or the generation of thousands of genomic markers (i.e., single-nucleotide polymorphisms (SNPs)) distributed throughout the genome via reduced-representation sequencing, sufficient to facilitate population-level estimation of metrics including diversity, relatedness, population structure, and introgression in an efficient, cost-effective manner (Ba et al., 2017; Chen et al., 2016; Peek et al., 2019; Rexer-Huber et al., 2019; Rick et al., 2019). Population-level reduced-representation sequencing (including restriction-enzyme associated DNA sequencing (RADseq; Baird et al., 2008), double-digest RADseq (ddRADseq; Peterson et al., 2012), and genotyping-by-sequencing (GBS; Elshire et al., 2011)) is an approach that can produce thousands of variant sites for high-resolution population genomic analyses (Davey et al., 2011; Davey & Blaxter, 2010; Narum et al., 2013) and as such has wide applicability for conservation (Andrews et al., 2016; Seabury et al., 2011; Wright et al., 2019). Unlike other reduced-representation approaches, GBS includes an amplification step, but fewer steps overall (Elshire et al., 2011). Large genomic marker sets are expected to provide greater power to resolve questions relating to hybridisation and introgression (similar to that observed when estimating genetic diversity and differentiation (Fischer et al., 2017) and relatedness (Galla et al., 2020)). However, studies comparing the enhanced resolution of genomic data with previous genetic analyses related to hybridisation in a conservation context are thus far limited (Table 1). Here we demonstrate the utility of genomic tools for assessment of the impacts of hybridisation in a critically endangered wading bird.

**Table 1.**
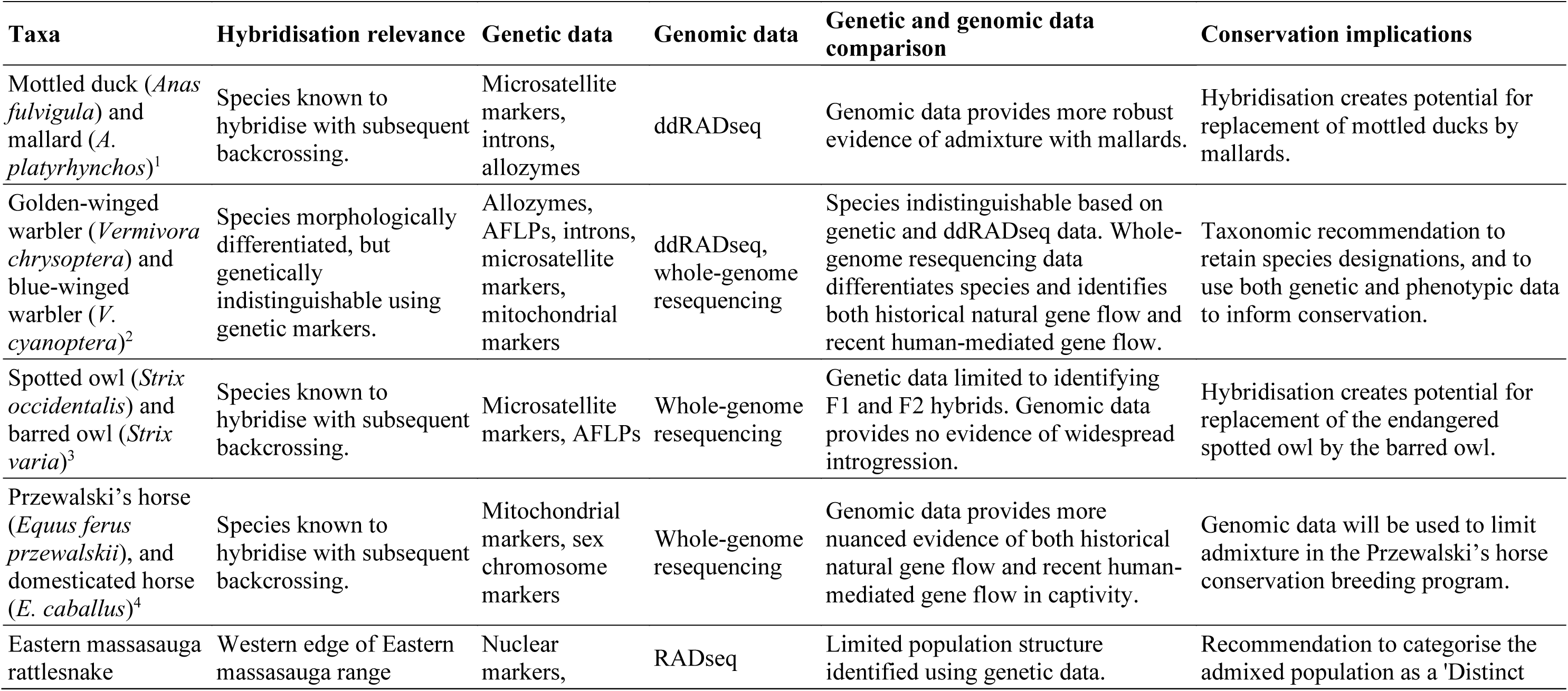
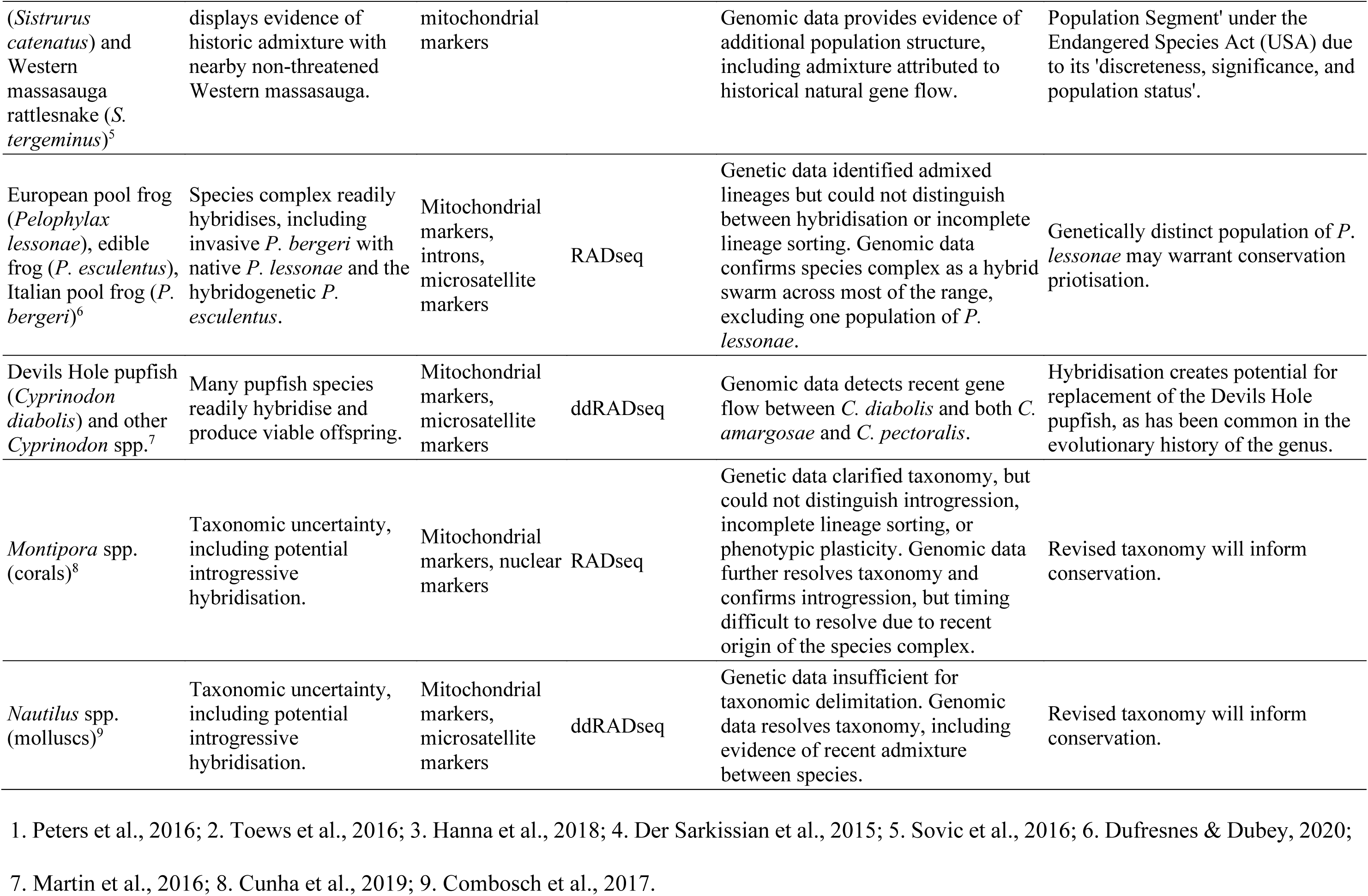
Interspecific hybridisation research in a conservation context where both genetic and genomic data have been generated and both have been used to assess hybrids or hybridisation. ‘Conservation relevance’ is used to mean that the hybridisation involves a threatened species, and the outcomes of the study have implications for conservation and/or present recommendations for conservation management. Web of Science search (19 July 2020): ALL FIELDS: (((genetic AND genomic) AND (hybridization OR introgression OR admixture) AND (conservation OR threatened OR endangered)) AND (fish OR bird OR mammal OR marsupial OR amphibian OR reptile OR plant OR invertebrate OR insect OR mollusc OR crustacean)). Of the 237 results, to capture empirical research that used high-throughput genomic sequencing techniques only, reviews and editorials, and articles published prior to 2014 were excluded. From the remaining 119 results, studies assessing intraspecific admixture or with no clear conservation relevance were excluded.

One of the world’s rarest bird species, the Aotearoa New Zealand kakī (black stilt, *Himantopus novaezelandiae*) provides a quintessential example of a threatened species affected by human-induced hybridisation (BirdLife International, 2018; Robertson et al., 2016). Anthropogenic impacts resulted in population decline during the 1900s, with numbers falling to approximately 23 individuals comprising a single population in Te Manahuna/the Mackenzie Basin in 1981 (Pierce, 1984b; Steeves et al., 2010). Adaptive conservation management of kakī, including predator control throughout Te Manahuna and a programme of captive breeding and rearing for translocation have been integral to increase the kakī population to 169 wild adults in 2020 (Hagen et al., 2011; Heezik et al., 2005; Keedwell et al., 2002; Maloney & Murray, 2001; Reed et al., 1993; Steeves et al., 2010).

Research to date based on pedigree and genomic data indicates that despite the population history of kakī, the population exhibits relatively low levels of inbreeding and relatedness among individuals (Galla, 2019; Galla et al., 2019). However, distorted allele frequencies in small populations can produce an apparent excess of heterozygotes, which would also result in underestimated levels of inbreeding, and such skewed results can be difficult to correctly assess in the absence of a comparable large, outbred population (Kardos et al., 2015). Inbreeding depression in kakī manifests reduced hatching success (Hagen et al., 2011), and so breeding has been carefully managed by the DOC’s Kakī Recovery Programme (summarised by Galla et al., 2020). Along with predation and altered habitat availability, interspecific hybridisation may pose a threat to species recovery (Steeves et al., 2010). The Aotearoa New Zealand population of congeneric Australian pied stilts (hereafter referred to as poaka; *Himantopus himantopus leucocephalus*) self-introduced from Australia at least 200 years ago, and anthropogenic impacts facilitated expansion of the species’ range (Pierce, 1984b). Limited mate choice and a male sex bias among kakī when kakī numbers were declining promoted hybridisation between kakī and poaka, producing viable, fertile hybrid offspring (Pierce, 1984a; Steeves et al., 2010).

Current kakī conservation management policies reflect the conservation value of non-admixed kakī, which are individuals with pure-black plumage that genetically assign to kakī based on a small set of mitochondrial and microsatellite markers (Maloney & Murray, 2001; Reed et al., 1993; Steeves et al., 2010). Steeves et al. (2010) confirmed the genetic distinctiveness of kakī, and found no evidence of introgression from poaka except in a single individual. This finding was attributed to outbreeding depression that negatively impacts fitness of female hybrid offspring, combined with an ephemeral skewed sex ratio and active management to exclude hybrids (Steeves et al., 2010). However, Steeves et al. (2010) acknowledged that the small marker set used may not be representative of genome-wide variation, and thus may lack the power to detect low levels of introgression resulting from hybridisation (see also Brumfield, 2010).

Here, we use population-level GBS and a reference-guided approach to SNP discovery for kakī, known hybrids, poaka, and Australian pied stilts. We infer the genomic extent and pattern of introgression due to hybridisation between kakī and poaka in the contemporary kakī population, as compared with that of previous genetic analyses. Determining the utility of genetic and genomic markers for detection of introgressive hybridisation is essential not only for the conservation of kakī, but also for detecting hybridisation in threatened species more broadly.

## 2 Materials and Methods

### 2.1 Sample collection and DNA extraction

Following Steeves et al. (2010), individuals sampled herein were grouped by plumage phenotype (‘node’) as categorised by Pierce (1984a) and used by the Department of Conservation’s Kakī Recovery Programme. Poaka and pied stilts (plumage nodes A–C2; Pierce, 1984a) were labelled ‘pied’, completely black node J individuals were labelled ‘kakī’, and individuals of intermediate plumage (nodes D1–I/J) or known hybrid parentage were labelled ‘hybrid’ (Supplementary Table 1). Extracted genomic DNA (gDNA) was available from 80 stilt samples including kakī, Australian pied stilts and poaka, and hybrids from previous genetic analyses.

We extracted gDNA from an additional 155 feather samples collected as part of regular handling practices for kakī under Aotearoa New Zealand’s Department of Conservation (DOC) ethics approvals (AEC #283) at the DOC’s Kakī Recovery Programme, Twizel, and the Isaac Conservation and Wildlife Trust kakī captive rearing facility, Christchurch, Aotearoa New Zealand. Pedigree information is recorded for all kakī individuals as part of routine Kakī Recovery Programme management, extending up to seven generations. In addition, blood samples were collected as part of routine health checks from two Australian pied stilts at Adelaide Zoo (provided under a Royal Zoological Society of South Australia Specimen Licence Agreement; Import Permit #2016061954), two node B poaka from Auckland Zoo (under Auckland Zoo animal ethics approval), along with a tissue sample from one poaka from Hawke’s Bay, North Island (Figure 1). Poaka from the North Island were preferentially sampled due to a low likelihood of recent contact with kakī, minimising the chance of these individuals having recent hybrid ancestry. We extracted gDNA from all samples using a Thermo Scientific™ MagJET™ Genomic DNA kit, following Protocol E (manual genomic DNA purification from up to 20 mg tissue). We quantified gDNA for all samples using a NanoDrop™ 8000 Spectrophotometer to assess viability for GBS. GBS guidelines from our sequencing provider (AgResearch Ltd., Mosgiel, Aotearoa New Zealand) recommended a total of 1 µg of DNA per sample at a concentration of 80–150 ng/μL, and 260/280 and 260/230 ratios of 1.80–2.00 and > 1.0 respectively.

**Figure 1.**
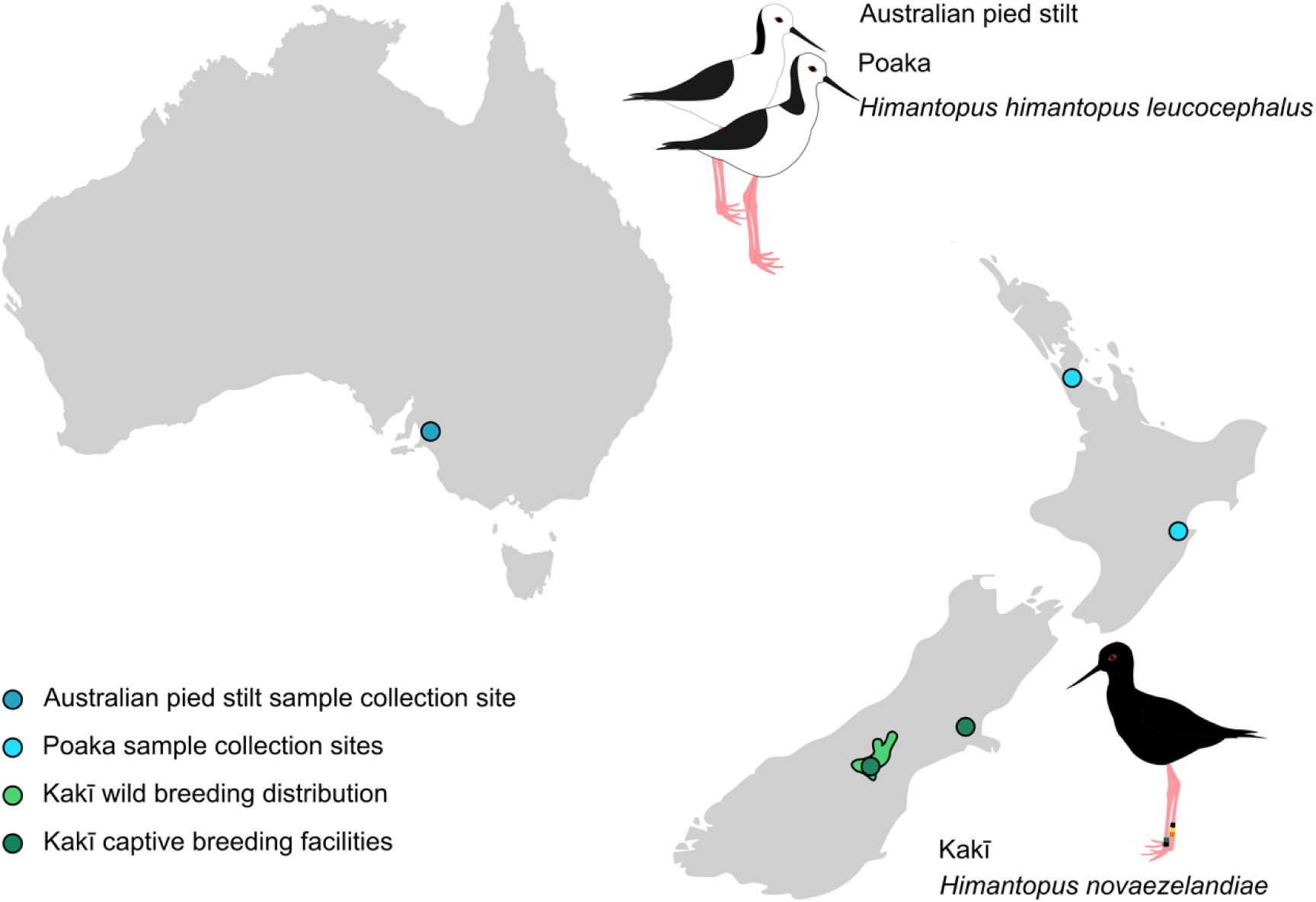
Locations of kakī captive breeding facilities and wild breeding distribution, and Australian pied stilt and poaka sampling sites in Australia and Aotearoa New Zealand (maps not to scale).

### 2.2 Genotyping-by-sequencing

We selected GBS as an appropriate reduced-representation sequencing method based on lower cost and availability within Aotearoa New Zealand (Galla et al. 2016). Of the available DNA extracts, 145 samples (130 kakī, six pied stilts, and nine hybrids) had DNA of sufficient quality and quantity as per GBS recommendations. This included 74 males and 62 females, with nine individuals of unknown sex (Supplementary Table 1). Nineteen individuals included in the previous genetic analysis by Steeves et al. (2010) had gDNA of sufficient quantity and quality for inclusion here (Supplementary Table 1). We prepared two 96-well plates of samples for GBS, containing 145 samples, two negative controls (DNA-free controls) per plate, and three positive controls (replicate samples) across plates. We diluted samples with gDNA concentration > 150 ng/μL to 100 ng/μL, and supplied ∼1 μg of DNA for all samples. Optimisation and sequencing of GBS was conducted by AgResearch Ltd. The optimised GBS protocol used a double-digest with restriction enzymes PstI-MspI. A single library was generated for all 145 samples and controls with a fragment length filter of 193– 500 bp including adapter sequences. This library was sequenced on one lane of Illumina HiSeq 2500 v4 sequencing for 101 cycles.

### 2.3 Reference-guided variant discovery

We assessed sequence quality with FastQC v0.11.5 (Andrews, 2010), and confirmed absence of contamination in negative controls through BLAST searches against the nucleotide database (Altschul et al., 1990). To provide accurate SNP discovery, we mapped GBS data to the kakī reference genome (Galla et al., 2019). We demultiplexed and filtered raw sequences with Sabre v1.0 (Joshi, 2013) and adapter trimmed with Cutadapt v1.17 (Martin, 2011). We indexed genomes and mapped the processed GBS reads to the genomes with BWA-MEM in BWA v0.7.17 (Li & Durbin, 2009). We pre-processed the mapped reads for variant discovery by adding read-group information with SAMtools, marking duplicates with Picard v2.18.0 (Picard Toolkit, 2019), and realigning indels with the Genome Analysis Toolkit (GATK) v3.5 (McKenna et al., 2010).

We compared five independent pipelines for reference-guided variant discovery. The ‘GATK’ pipeline used GATK’s HaplotypeCaller and GenotypeGVCFs to call variants. The ‘Samtools’ pipeline used SAMtools v1.7 mpileup and BCFtools v1.6 variant caller (Li, 2011). The ‘Platypus’ pipeline used the callVariants tool in Platypus v0.8.1 (Rimmer et al., 2014) with minimum mapping quality of 20, minimum base quality of 20, minimum depth to call a variant of 2, and flag to generate indels set. The ‘Stacks’ pipeline implemented Stacks v2.2 (Catchen et al., 2013) reference-guided pipeline with default parameters. The mapped sequence reads were passed as input to GATK, Samtools, Platypus, and Stacks. The fifth pipeline, ‘Tassel’, was run independently with the raw multiplexed GBS data passed as input to TASSEL5-GBS2 v5.2.39 (Glaubitz et al., 2014). Tags were extracted from the data set with a minimum quality score of 10, and then passed to BWA v0.7.12 for alignment against the reference genome. The resulting SAM file was passed back to TASSEL5-GBS2 for variant discovery with default settings, and all SNPs with quality ≥ 10 were retained.

### 2.4 Variant processing

By using a reference-guided approach, genomic location data was available for all variants, and so we could compare variants produced across the five pipelines using VCFtools v0.1.15 (Danecek et al., 2011) function *vcf-compare* following standardisation of variant call format files with *vcf-convert*. We visualised the intersections of common variants among pipelines with the package UpSetR (Conway et al., 2017; Lex et al., 2014) implemented in R v3.5.1 (R Core Team, 2018). To improve confidence that the SNPs discovered were true SNPs rather than the result of sequencing or mapping error, we produced a single variant set comprising all variants detected via at least three pipelines from the intersections of variants common to multiple pipelines generated using *vcf-isec*, *vcf-merge* and *vcf-sort* functions in VCFtools. To produce a set of biallelic SNPs to investigate admixture between kakī and poaka, we removed indels and multiallelic SNPs from the composite variant set using VCFtools. To confirm absence of contamination and replicability of lab processes, preliminary filtering tests and downstream analyses had retained negative and positive controls, and once confirmed, these controls were removed. We then excluded sites with > 10% missing data, a minor allele frequency < 0.01, and a minimum quality score < 20. SNPs with mean depth over all individuals between 5× and 200× were retained, and individuals with > 50% missing data across all sites were excluded (final n = 140). Due to the putative nature of the sex chromosomes in the kakī reference genome, preliminary analyses were conducted both including and excluding SNPs located on putative sex chromosomes. In addition, we subsampled the individuals to include only a single individual from known family groups based on pedigree data, and excluded known hybrid offspring. No differences in the extent of introgression was observed from downstream analyses for any of these approaches (data not shown), but we conservatively excluded SNPs located on putative sex chromosomes.

### 2.5 Discriminant Analysis of Principal Components

In an exploratory multivariate approach to population clustering, we conducted Discriminant Analysis of Principal Components (DAPC) with adegenet v2.1.1 (Jombart, 2008; Jombart et al., 2010; Jombart & Ahmed, 2011; Jombart & Collins, 2015b) in R v3.5.1. DAPC attempts to partition variance in a between-group and within-group manner to maximise the discrimination between groups. Using a multivariate approach allows for fine-scale assessment of population structure, without relying on population genetic models, and so is independent of the assumptions of HWE or linkage equilibrium associated with population structuring analyses (Jombart et al., 2010). DAPC uses *a priori* information of the number of clusters present in the data set (two, kakī and pied stilts including poaka), and then assesses the discriminants that best explain those clusters. To prevent overfitting of the data, we optimised DAPC parameters using the Bayesian Information Criterion (BIC) and *a*-scores, and performed cross-validation following Jombart & Collins, 2015a). The *a*-score measures the trade-off between the power of discrimination and potential for overfitting the data, using a randomisation of the data to determine when cluster assignment is successful due to the analysis or due to random discrimination, and penalises the reassignment score by the number of retained principal components (PCs). Cross-validation confirmed the appropriate numbers of PCs, using a random seed to produce 1,000 replicate runs with a training set of 80% of the data across up to sixty PCs. The accuracy of the retained PCs was then tested with the remaining 20% of the data, and the PCs retained for the final DAPC were based on those which produced the lowest mean squared error and highest mean success. The optimised DAPC analysis was visualised to infer species differentiation and individual clustering. We also implemented PCA and additional clustering analyses with the package SNPRelate v1.20.1 in R v3.6.3 for comparison with DAPC and ADMIXTURE analyses (Zheng et al., 2012). Results produced with SNPRelate were concordant with those of DAPC and ADMIXTURE (Supplementary File 1).

### 2.6 Analysis of introgression with ADMIXTURE

To estimate individual assignment to population clusters and to detect introgression, we analysed each SNP set with a maximum likelihood method implemented in ADMIXTURE v1.3.0 (Alexander et al., 2009). To minimise stochasticity across multiple runs, we conducted 100 iterations of ADMIXTURE analysis with each SNP set for *K* = 1–6, where *K* represents the hypothesised number of population clusters, using a random seed, ten-fold CV, and with point estimation terminating when the change in log-likelihood increased by < 0.0001. The range of *K*-values was selected independently of the results of DAPC analysis, allowing for differentiation between the two species (kakī and pied stilts), along with potential population structuring among kakī, or differentiation between Australian pied stilts and poaka. To determine the most appropriate value of *K* for each SNP set, we averaged CV error across the 100 iterations and visualised the results, with the lowest CV error representing the most likely *K*.

There are two potential introgression scenarios that are of particular concern for kakī conservation managers. In the first scenario, a substantial amount of introgression would be identified among a small number of kakī due to recent undetected hybridisation in the wild. Accurate identification of any such individuals will inform breeding management decisions where the goal is to avoid genetic mixing between kakī and poaka. In addition, changes in wild population management may be required to detect such rare hybridisation events. In the second scenario, a moderate amount of introgression previously undetected with genetic markers may be observed across a greater number of kakī, resulting from the presence of a small number of dark hybrids among the founding individuals in the captive breeding programme (Galla et al., 2020), which may have been exacerbated by subsequent selection maintaining introgressed genes. In this second scenario, breeding management practices may be altered to take into account the proportion of introgression in individuals to minimise genetic admixture. In both scenarios, robust and accurate detection of introgression is requiredto determine future research and management needs. Thus we defined a stringent threshold of > 0.95 individual probability to assign individuals as kakī, informed by the strong genetic differentiation between these two species (Steeves et al., 2010).

We visualised mean assignment probabilities (*Q*-values) across all iterations with pophelper v2.3.0 (Francis, 2017) in R v3.5.1. We used pophelper for file conversion for input to CLUMPP to handle label switching. Consensus *Q*-values for each individual were calculated with the Greedy algorithm over 100 iterations in CLUMPP vMacOSX 1.1.2 (Jakobsson & Rosenberg, 2007), and we visualised the results with pophelper in R v3.5.1. We manually assessed the final *Q*-matrix for all individuals using the predefined assignment threshold to assign individuals as kakī. To reduce any potential influence of related individuals from the ADMIXTURE analysis, the SNP set was filtered to exclude known related kakī individuals based on pedigree data (a single individual was retained among each known family group). This SNP set also excluded poaka to avoid any influence of potential admixture from kakī among those individuals. ADMIXTURE and DAPC analyses were conducted as above, and revealed no differences in population structure or patterns arising from family relatedness or inclusion of poaka (Supplementary Figures S1 and S2).

### 2.7 Combining pedigree data with genomic population assignment data

Following all admixture and population clustering analyses, only six node J individuals were identified with < 1.00 kakī assignment. A small number of hybrid individuals are included among the founders in the kakī pedigree (Galla et al., 2020). To determine whether these assignment probabilities could be attributed to known hybrid ancestry (< 7 generations deep), we used the kakī pedigree (Galla et al., 2020) to assess the ancestry of these six individuals.

## 3 Results

Following a reference-guided multi-pipeline approach to SNP discovery and filtering for 145 stilts, a total of 140,948 SNPs were used in downstream analyses which detected no evidence of introgression from poaka into kakī. Of the 250 gDNA samples available, 145 extractions contained DNA of the required quantity and quality for GBS, including 66 of the 106 (63.2%) adults alive in the wild kakī population when this study began in 2017. GBS of the pooled set of kakī, Australian pied stilts and poaka, and interspecific hybrids produced a total of 303,639,199 raw sequences with length 35–101 bp and high sequence quality. Demultiplexing produced an average of 2,024,530 ± SD 1,031,208.21 reads per sample (Supplementary Table 2), and no samples failed to sequence. Negative controls produced a low number of reads (mean = 2,585 ± SD 1554.76 reads per negative). Contamination checks of negative controls produced no matches to the BLAST nucleotide database.

### 3.1 Variant discovery and filtering

Mapping of trimmed, filtered reads for all samples to the reference kakī genome produced an average of 1,138,306.05 ± SD 597,406.95 mapped reads per individual (Table 2). This represents an average of 85.4% reads per sample successfully mapped to the kakī reference genome.

**Table 2.** Sequencing outputs and mapping success of GBS data from kakī, Australian pied stilts/poaka, and interspecific hybrids averaged by species. Overall includes all samples along with negative and positive controls.

|  | Mean reads | Mean cleaned reads | Mean reads mapped to kakī genome |
| --- | --- | --- | --- |
| Kakī (n = 130) | 2,094,197.47 | 1,378,947.26 | 1,211,024.32 |
| Pied stilts (n = 6) | 1,722,470.33 | 966,686.33 | 879,153.17 |
| Hybrids (n = 9) | 1,181,117.00 | 812,732.00 | 732,418.11 |
| Overall | 1,971,321.13 | 1,293,796.12 | 1,138,306.05 |

Variant calling with GATK produced the fewest variants (35,441), while SAMtools produced the most (488,940, Fig. 2). There were 177,437 variants common to ≥ 3 pipelines (Fig. 2). Despite GATK producing the fewest variants among the five pipelines, the majority (92.68%) were retained in the common variant set. SAMtools had the lowest proportion (35.14%) of discovered variants retained in the common set. A total of 15,851 SNPs remained after filtering. The five individuals that had produced the fewest raw sequences (26,725–313,884 reads) were subsequently excluded due to high (> 50%) levels of missing data. The total genotyping rate was 96.91%.

**Figure 2.**
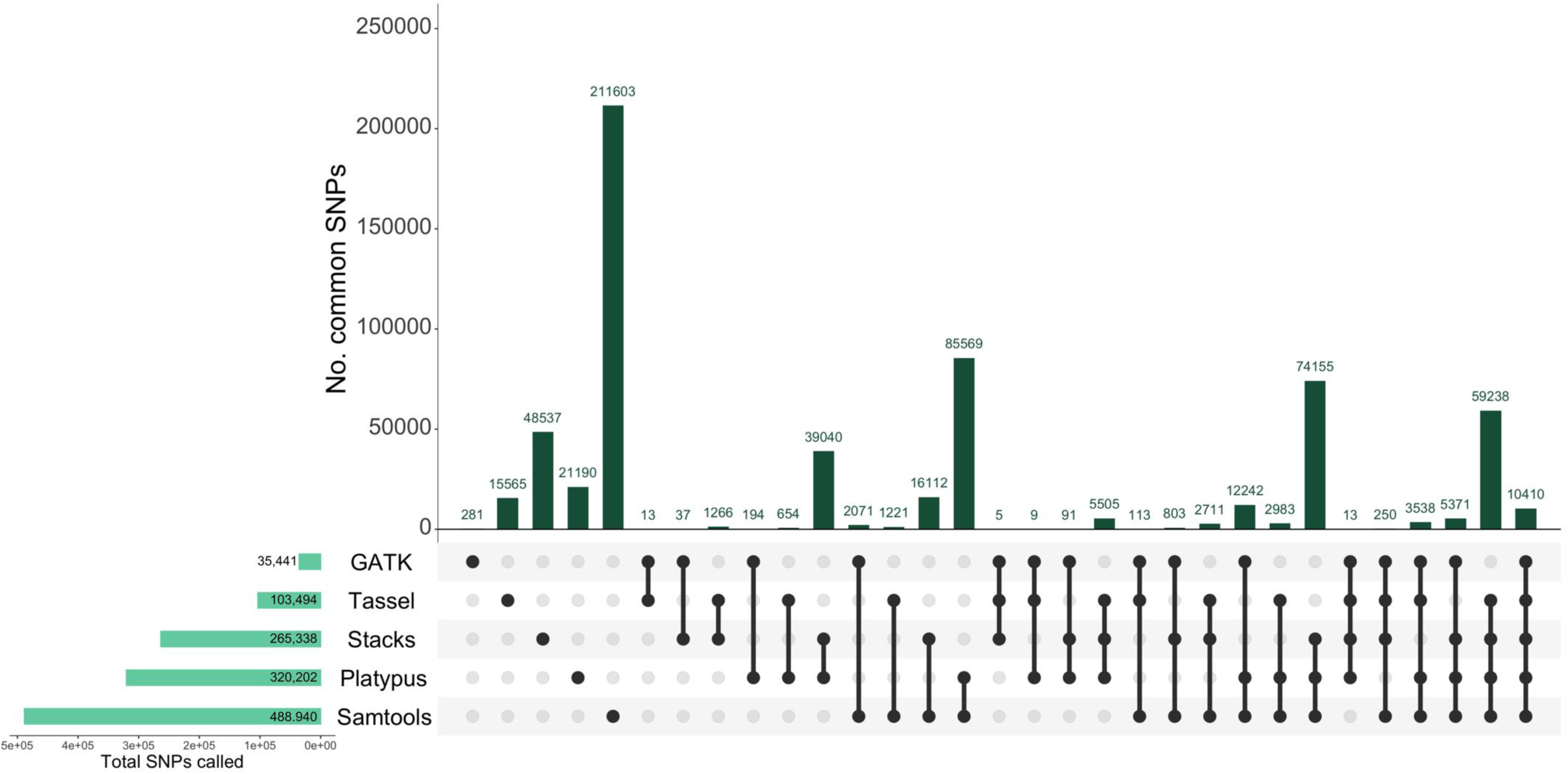
UpSetR plot of the intersections of the total variants discovered from GBS data for kakī, Australian pied stilts and poaka, and interspecific hybrids across five variant discovery pipelines: GATK, Platypus, Samtools, Stacks, and Tassel. Bottom left bars represent the total number of variants discovered with each pipeline, while the main bar plot represents the number of variants common to multiple pipelines as indicated by the filled circles below.

Exploratory statistics indicated an even distribution of SNPs throughout the genome, with a mean density of 0.014 ± SD 0.158 SNPs/kb (Table 3). Mean depth of coverage per individual was 11.663×. Kakī were well-differentiated from pied stilts with mean F_ST_ = 0.622 (Table 3). Per-population summary statistics identified pied stilts as having higher diversity in terms of nucleotide divergence, a greater number of variant sites, and more private alleles than either kakī or hybrids (Table 4). Kakī displayed the lowest nucleotide diversity and fewest polymorphic sites among the predefined groups (Table 4).

**Table 3.** Summary statistics for single-nucleotide polymorphisms (SNPs) produced from GBS data for stilts. Mean ± standard deviation (SD). KB = kilobase, HWE = Hardy-Weinberg Equilibrium, FDR = False Discovery Rate, F_ST_ = measure of population differentiation, Ts/Tv = ratio of transitions to transversions.

|  | Stilt SNP metrics |
| --- | --- |
| Total SNPs | 15,851 |
| Total samples | 140 |
| Total Kakī / Pied / Hybrid individuals | 125 / 6 / 9 |
| Mean depth per SNP per individual | 11.663 ± 6.107 |
| Mean per SNP depth | 1,632.77 ± 668.176 |
| Mean SNP quality | 1,248.19 ± 2447.43 |
| Mean frequency of missing data per individual | 0.031 ± 0.073 |
| Mean SNPs/KB | 0.014 ± 0.158 |
| Total singletons/private doubletons | 0 |
| Mean nucleotide diversity; $\pi$ | 0.0912 ± 0.100 |
| SNPs deviating from HWE (FDR-corrected $P \leq 0.05$) | 758 |
| Ts/Tv ratio | 3.894 |
| Weir & Cockerham mean $F_{ST}$ (Kakī v Pied) | 0.622 |
| Weir & Cockerham weighted mean $F_{ST}$ (Kakī v Pied) | 0.637 |

**Table 4.** Population summary statistics from the three single-nucleotide polymorphism (SNP) sets produced from GBS data for kakī, Australian pied stilts/poaka, and interspecific hybrids, as calculated during format conversion from VCF to PLINK.

|  | Kakī | Pied | Hybrid |
| --- | --- | --- | --- |
| Mean samples per locus | 121.990 | 5.542 | 8.141 |
| Polymorphic sites | 6,729 | 14,699 | 13,621 |
| Private alleles | 389 | 1,100 | 122 |
| Mean nucleotide diversity ($\pi$) | 0.057 | 0.396 | 0.235 |

### 3.2 Discriminant Analysis of Principal Components

DAPC analysis identified Australian pied stilts and poaka as clustering separately from kakī, with hybrids intermediate to the two species, though grouping more closely with kakī than poaka (Fig. 3). Two node I/J hybrid individuals (DNA777 and DNA779) were found to cluster with kakī. These results were concordant with clustering analyses conducted with SNPRelate (Supplementary File 1)

**Figure 3.**
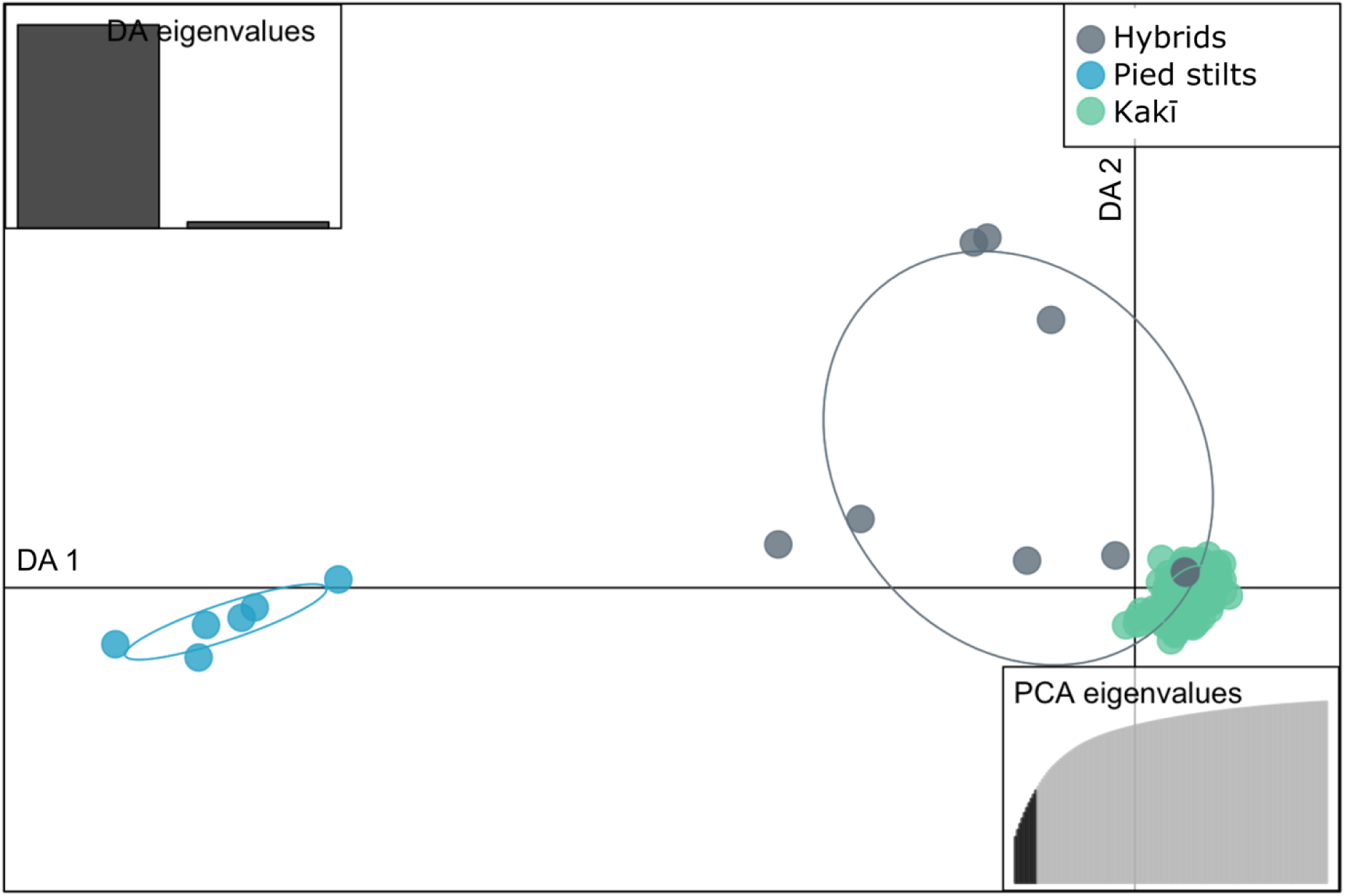
Scatterplot of a Discriminant Analysis of Principal Components (DAPC) produced from GBS data for kakī, Australian pied stilts and poaka, and interspecific hybrids conducted in adegenet. Individual points are coloured according to plumage node and/or known ancestry as one of: kakī (*Himantopus novaezelandiae*), Australian pied stilts/poaka (*H. himantopus leucocephalus*), or interspecific hybrids. The closer individuals are to one another in the plot, the more likely they are to have shared genetic ancestry. DAPC analysis was optimised using *a*-score, cross-validation, and BIC to derive the appropriate number of principal components (2 discriminants, 10 PCs). The 67% inertial ellipses around each cluster represent the variance of the clusters depicted. The insert of PCA eigenvalues represents the variation explained by the PCs, and the insert of DA eigenvalues represents the magnitude of variation explained by the two discriminants.

### 3.3 ADMIXTURE analysis of introgression

*K* = 2 was indicated as the most likely number of clusters for ADMIXTURE analysis based on CV error values, consistent with previous genetic results confirming taxonomic delimitation (Steeves et al., 2010), and therefore only results for *K* = 2 for are reproduced here. All individuals categorised as kakī (node J individuals) had assignment probabilities to the kakī cluster above the pre-defined 0.95 threshold, and only six individuals were assigned as kakī with probability < 1.00 (kakī mean *Q* = 0.9992 ± SE 0.0009; Fig. 4, Table 5, Supplementary Table 3). Both Australian pied stilts and the poaka individual Poaka1 were assigned with 1.00 probability to the pied stilt cluster. Assignment probabilities to the kakī cluster for hybrids ranged from 0.2301 (DNA2113) to 1.00 (DNA777 and DNA779; hybrid mean *Q* = 0.6399 ± SD 0.0175; Supplementary Table 3).

**Figure 4.**
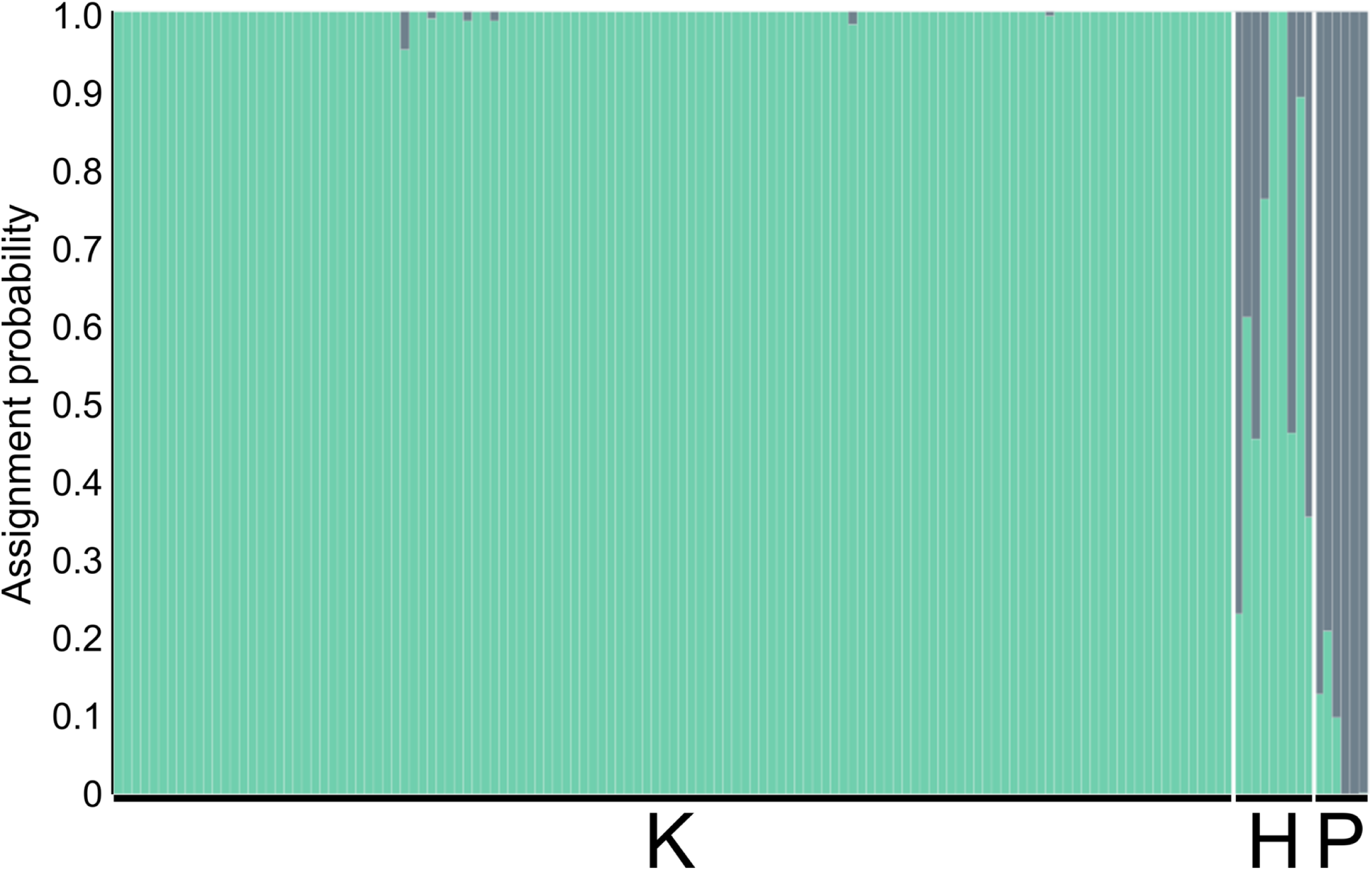
Assignment probabilities for kakī (K), Australian pied stilts and poaka (P), and interspecific hybrids (H) produced via pophelper visualisation of CLUMPP-permuted ADMIXTURE results when *K* = 2. Each individual is represented by a vertical bar, with colours indicating the assignment probability to the kakī (green) or Australian pied stilts/poaka (grey) cluster.

**Table 5.** Mean individual assignment probabilities (*Q*-values) produced through 100 iterations of ADMIXTURE analysis for kakī (node J individuals with known node J ancestry) that were assigned to the kakī cluster with probabilities above the 0.95 threshold, but below 1.00.

| DNA ID | Mean assignment probability, $Q$ |
| --- | --- |
| DNA1252 | 0.9514 |
| DNA1429 | 0.9838 |
| DNA1620 | 0.9882 |
| DNA1694 | 0.9945 |
| DNA897 | 0.9882 |
| DNA932 | 0.9910 |

### 3.4 Combining pedigree and genomic population assignment data

All node J individuals were assigned as kakī with probabilities above the 0.95 threshold. Pedigree assessment revealed only 17 kakī individuals included in this study had no recorded hybrid ancestry (i.e., all of their recorded ancestors were individuals with plumage node J representing completely black birds). Among the node J kakī individuals identified as having 0.9500–0.9999 probability of assignment to the kakī cluster, all individuals had at least one node I or I/J individual in their recorded ancestry. Individual DNA897 had the most recent hybrid ancestry, with an I/J ancestor three generations deep. Individual DNA1252 had the lowest probability of assignment to the kakī cluster among all kakī (Table 5), and while this individual did not have more frequent or more recent recorded hybrid ancestry than other kakī here, there is no recorded ancestry for the paternal lineage. The mother of individual DNA1252 was included in sequencing and analyses, and was not identified as having any hybrid ancestry, suggesting a paternal lineage origin for the relatively lower probability of assignment as kakī. Individual DNA1694 has a relatively deep pedigree among kaki, spanning seven generations. This individual also has the most frequent incidence of recorded hybrid ancestry among these individuals, with a node I/J ancestor four generations deep, three further I/J ancestors six generations deep, and a node I ancestor seven generations deep. The two hybrid individuals that were consistently assigned to the kakī cluster were siblings DNA777 and DNA779, with a node I/J mother, and a node I/J ancestor three generations deep.

## 4 Discussion

With the development of genomic sequencing and associated methodologies, conservation management of threatened species can now incorporate high-resolution genomic tools to more robustly explore threats to species recovery, such as anthropogenic hybridisation (Allendorf et al., 2010; Avise, 2010; Ekblom & Galindo, 2011; Gompert, 2012; Primmer, 2009). As discussed by Allendorf et al., (2001), hybridisation of a threatened species with a more common non-native species can cause widespread introgression and the formation of a hybrid swarm replacing the distinct species. Studies to date indicate that while using a small number of genetic markers can provide important baseline data for conservation management (e.g., genetic sexing, (Steeves et al., 2010); brood parasitism, (Overbeek et al., 2017; Overbeek et al., 2020), such marker sets have proved less robust than genomic markers for analyses of parentage (Tokarska et al., 2009), relatedness (Galla et al., 2020), intraspecific population structure (McCartney-Melstad et al., 2018), and introgression (Parejo et al., 2018).

### 4.1 Impacts of hybridisation in stilts

All 130 kakī genotyped here—representing 63% of the contemporary adult population at the time of sampling—were assigned as kakī with > 0.95 probability. These data confirm that despite hybridisation with poaka, genetic admixture into kakī has not occurred, concordant with earlier genetic analysis (Steeves et al., 2010). These results contrast with those of the study by Toews et al. (2016) in which analysis of GBS data could not distinguish between two hybridising species of warblers, and whole-genome sequencing data was required. This is likely due to the very low differentiation observed between the two warbler species, compared with the strong differentiation between kakī and pied stilts.

No individuals with pure-black plumage (node J) were identified as cryptic hybrids, with all individuals assigned to the kakī cluster with probabilities above the predefined 0.95 threshold. Among the sampled individuals, 29 kakī had at least one hybrid ancestor three generations in the past, and these ancestors were all dark hybrids (node I or I/J). No kakī individuals included in this study had recorded hybrid ancestors with plumage lighter than node I. These results indicate that with every generation of kakī backcrossing following hybridisation, the proportion of introgressed genes is increasingly likely to be replaced by kakī genetic material. Indeed, two individuals identified as hybrids (DNA777 and DNA779) with a node I/J mother were assigned as kakī with probability of 1.00 across all analyses. This may indicate that even a small number of generations of backcrossing with kakī can rapidly overwhelm introgression from poaka. The current kakī recovery strategy excludes any non-node J birds from active management (i.e., excluding any offspring of node I/J birds from the captive rearing programme; Steeves et al., 2010).

The finding of no introgression from poaka to kakī likely results from a combination of factors. First, the management strategy enacted by the DOC’s Kakī Recovery Programme to avoid genetic admixture of kakī has successfully minimised opportunities for hybridisation between these species. This management has been responsive to the results of genetic analysis that led to the exclusion of all non-node J individuals from conservation management (Steeves et al., 2010). Intensive population monitoring of breeding pairs and assessment of putative hybrids using the microsatellite panel has enabled practitioners to break up mixed pairs (allowing kakī to re-pair with kakī) and exclude hybrids from the captive breeding programme (Maloney & Murray, 2001). Ongoing kakī recovery has resulted in a relatively balanced sex ratio in the wild, and combined with the strong positive assortative mating of kakī, has minimised the likelihood of kakī breeding with non-kakī (Steeves et al., 2010). Moderate outbreeding depression and stochastic processes have also contributed to reduce the reproductive success of hybrids (Steeves et al., 2010), further limiting the likelihood of introgressed material to be maintained in the population.

### 4.2 Implications for kakī conservation

The results presented here provide evidence that active conservation management designed to minimise hybridisation can be effective in maintaining species integrity, and support the ongoing management strategy of poaka and hybrid exclusion from the captive management programme based on the results of genetic analysis (Steeves et al., 2010). The goals of current kakī management to avoid genetic admixture are appropriate due to the strong differentiation between species observed here, reduced fitness of hybrid offspring, and the previous observation of moderate levels of genetic diversity among kakī compared with other threatened Aotearoa New Zealand endemic birds (Steeves et al., 2010). Despite the increased resolution of genomic data, when individuals with anomalous plumage are observed in the captive breeding and rearing facility, microsatellite genotyping remains the most cost- and time-efficient, low-complexity method for confirming species status (Table 6; Overbeek et al., 2017; Overbeek et al., 2020), and this may be the case for other species threatened by hybridisation. Early confirmation of hybrid status with genetic tools is key, as kakī do not become fully black until about two years of age, although the Kakī Recovery Team are experienced at identifying putative hybrid offspring appearing as unusually ‘pale’ chicks/juveniles. Evidence of reduced fitness of hybrid offspring (Steeves et al., 2010) suggests avoidance of genetic admixture will continue to be prioritised over any potential gain in genetic diversity that may be facilitated through managed introgression (e.g., the inclusion of cryptic hybrids). Further, the absence of cryptic hybrids in the contemporary kakī population indicates continued avoidance of genetic admixture is possible for kakī (Steeves et al., 2010; this study). One key challenge of hybridisation to conservation is consideration of the impacts of inbreeding versus outbreeding (Liddell et al., 2021). Further research pertaining to inbreeding and inbreeding depression could enhance understanding of the consequences of maintaining these species as distinct entities, with particular focus on the impacts of population decline and ongoing small population size on kakī genetic diversity as compared with known impacts of outbreeding (Steeves et al., 2010). Nevertheless, the mitigation of risks associated with inbreeding by the DOC’s Kakī Recovery Programme, as compared with the challenges of managing similar risks associated with outbreeding mean the current management strategy remains the most appropriate.

**Table 6.** Comparison of the costs and benefits associated with a genetic approach (i.e., microsatellite panel) for assessing hybridisation in kakī with a genomics approach (i.e., genotyping-by-sequencing). With both platforms already established for kakī, cost per sample for additional small number of samples assessed each year is the primary deciding factor. All cost estimates are in British pounds (GBP) based on cost estimates in Aotearoa New Zealand.

|  | **Microsatellite panel** | **Genotyping-by-sequencing** |
| --- | --- | --- |
| Platform development cost | < 5,000 GBP. Includes development and screening of ~20 polymorphic loci, and genotyping of up to 94 individuals (based on a known decrease in costs since the estimate of Galla et al. (2016)). | ~3,000 GBP. Includes testing of restriction enzymes and adapter barcoding, plus DNA extraction and sequencing of up to 94 samples, with expected output of 30–60K SNPs discovered via a de novo approach to SNP discovery. |
| Platform development time | 3–4 months | 3–4 months |
| Cost per sample for additional samples | 10 GBP. Includes DNA extraction and quantification, and microsatellite genotyping at eight optimised loci. Excludes associated person-hours. | 30 GBP. Includes DNA extraction and quantification, and GBS. Excludes associated person-hours. |
| Lab time for additional samples | 1 week | 1–3 months |
| Analysis time | 1 week (including manual genotyping) | 4 weeks |
| Analysis requirements | Access to a standard desktop computer.<br>Access to allele-calling software (e.g., GeneMarker v2.2).<br>Access to and experience with genetic population clustering tools optional. | Access to a high-capacity computing system or capable computing cluster.<br>Access to and experience with a variety of population genomic and bioinformatic tools. |
| Additional benefits | Previous uncertainty regarding how representative of genome-wide patterns of introgression these data are has been resolved for kakī by comparison with GBS data, indicating the eight loci are sufficient for robust identification of non-kakī individuals.<br>All wet-lab work and analysis for additional samples can be run in-house. | Increased confidence in detection of introgression with large genome-wide SNP set compared with microsatellites.<br>No ascertainment bias associated with marker generation.<br>Once these data are generated, they can be implemented for a variety of downstream uses.<br>Potential for these data to have additional application in the future with genomic analysis developments. |
| Limitations | Caveats associated with how the microsatellite loci are generated, and what they were developed for (e.g., active avoidance of regions under selection, ascertainment bias from selecting the most heterozygous markers). This limits the type of data produced, and thus the types of analyses that can be performed.<br>Potential for human error associated with manual allele-calling, but mitigated with experience. | Additional sampling requires the SNP discovery pipeline to be run with all previous samples included every time, and so analysis does not become more efficient over time.<br>Potential for batch effects between different sequencing batches (Leigh et al., 2018).<br>Analyses limited by software and models available in a developing field. |

Under optimal circumstances, kakī recovery will continue, leading to increased numbers of kakī within Te Manahuna in the short-term, and the potential for natural expansion beyond the basin in the long-term. In the short-term recovery scenario, the success of the conservation breeding and rearing programme to date may mean that active management of the species could be scaled back. This may see a reduction in the number of adult kakī maintained in captivity for breeding, although management of wild nests, including egg-collection, artificial incubation and captive rearing are likely to continue to maximise population growth while kakī remain critically endangered. As kakī are one of the few threatened Aotearoa New Zealand birds to have maintained a population on the mainland despite the presence of invasive predators, and are capable of travelling long distances, active translocations are unlikely to be necessary to support natural expansion beyond Te Manahuna. In addition, it is unlikely that active management of any such expansion would be feasible. As such, management to minimise the likelihood of hybridisation within Te Manahuna will continue, but with the wide distribution of poaka across the country, future expansion into areas with high poaka densities may result in the increased prevalence of hybridisation that may promote genetic admixture. Therefore, avoiding hybridisation in the source population within Te Manahuna should be of high priority for conservation.

### 4.3 Impacts of hybridisation on poaka

The identification of poaka with pied stilt assignment probabilities < 0.95 may be a result of initial small population size on arrival to Aotearoa New Zealand, and subsequent hybridisation with kakī prior to species decline. Kakī only occur as vagrants in the North Island of Aotearoa New Zealand, observed in very low numbers since at least the 1950s (Pierce, 1984b). Given the limited contact between kakī and poaka in the North Island in recent years, we expected the poaka samples sourced from the North Island to produce assignment probabilities similar to those of the Australian pied stilts. However, both individuals sourced from Auckland Zoo were assigned to the pied stilt cluster with < 0.95 probability. The only North Island individual with an assignment probability comparable to those of the Australian pied stilts was the individual from Hawke’s Bay (Poaka1). This suggests that hybridisation early in the establishment of poaka may have resulted in introgression of kakī genetic material into an initially small poaka population that was not frequently supplemented by a substantial number of new immigrants, with introgressed material maintained in the expanding population despite subsequent backcrossing. Kakī introgression into poaka is supported by the observation of node A poaka having tarsal lengths outside the range observed among Australian pied stilts with no history of hybridisation, and poaka presenting a greater proportion of black plumage than is typical among Australian pied stilts (Pierce, 1984a).

### 4.4 Comparison of genetic and genomic approaches to introgression analysis

Reduced-representation sequencing approaches have proven to be efficient, robust, and cost-effective for variant discovery (Andrews et al., 2016; Davey et al., 2011; Elshire et al., 2011; Peterson et al., 2012). Here GBS was used as a relatively cost-effective approach to population-level genomic sequencing of non-model species, producing a set of species-discriminating SNPs. Initial development of a GBS system is markedly less expensive than development of a microsatellite panel (3000 GBP for GBS development using 94 samples in this study in 2018 compared with 5000 GBP development and testing of a microsatellite panel of approximately ten loci using 94 samples based on the estimate of Galla et al. (2016; Table 6). However, ongoing costs of the microsatellite panel per sample remain considerably lower than that of GBS (10 GBP/sample for a microsatellite panel compared with 30 GBP/sample for GBS; Table 6), and the time required from individual sampling to completion of analysis is substantially reduced. There are also fewer barriers to analysing microsatellite data (e.g., microsatellite genotyping and analyses can be conducted on a standard desktop computing system compared with the requirement of a high-performance computing system with access to and experience with a variety of bioinformatic tools necessary to analyse GBS data, Table 6). While the ability to more readily characterise genome-wide variation will make a genomics approach desirable for many conservation projects, the associated costs may limit uptake, especially when providing data for time-dependent decisions. Despite the increasing uptake of genomics approaches to answer questions pertinent to conservation management (Galla et al., 2016), the current greater costs and other transitional challenges (e.g., bioinformatic expertise) will likely maintain the conservation genomics gap for at least some species for the foreseeable future.

For kakī, the nature of GBS as a reduced-representation approach means that despite the large increase in data compared with the eight microsatellite loci used previously, this only represents < 1% of the 1.1 Gb kakī genome (Galla et al., 2019). While whole-genome resequencing on the scale achieved here was prohibitively expensive when this study was initiated, declining costs of resequencing have already overtaken reduced-representation sequencing. For example, combined with the kakī pedigree, whole-genome resequencing data is now being used to inform the kakī conservation breeding programme (Galla et al., 2020). Thus, should renewed population decline and a skewed sex bias cause hybridisation to increase and threaten kakī recovery in the future, or should the conservation value of cryptic hybrids be reconsidered, then a comparison of GBS with whole-genome resequencing data may be useful.

## 5 Conclusions

Studies comparing the utility of genetic and genomic approaches for generating estimates of population genetic diversity and differentiation indicate that large SNP sets generally outperform the small microsatellite sets typically used in conservation genetic studies (Hauser et al., 2011; Hohenlohe et al., 2013; Santure et al., 2010; Weinman et al., 2015). Thus, re-examining the extent of introgression between critically endangered kakī and non-threatened congeneric poaka using a genomic approach was essential to ascertain the efficacy of conservation management aimed at avoiding genetic admixture of kakī. While results are concordant between genetic and genomic approaches for kakī and poaka, this may not be the case for other species, particularly when hybridisation may be widespread (e.g., hybridisation between koloa maoli/Hawaiian duck (*Anas wyvilliana*) and the invasive mallard (*A. platyrhynchos*; Wells et al., 2019). Thus, we recommend that when genetic assessment has not been conducted, or there is uncertainty as to whether genetic data have adequately captured the impact of hybridisation, a genomic approach should be used. Further, we suggest that when genetic and genomic results are concordant – which we anticipate will be more likely for well-differentiated species – conservation managers can confidently continue to use genetic tools, particularly when these remain more efficient and cost-effective.

## 6 Data accessibility

Kakī are a taonga (treasured) species for Māori (the Indigenous people of Aotearoa New Zealand) and as such, genomic data derived from kakī are also recognised as taonga in their own right. Due to the tapu (sacred) nature of these data, the data presented here are hosted on a password-protected database at http://www.ucconsert.org/data/ and will be made available at the discretion of the kaitiaki of the iwi (tribes) and hapū (subtribes) associated with kakī. These data include raw genotyping-by-sequencing data, and the VCF comprising the unfiltered SNP set. Code for the analyses described is available on GitHub at https://github.com/natforsdick/Himantopus.

## 7 Acknowledgements

We extend our thanks to the tangata whenua who are kaitiaki for kakī, namely Te Rūnanga o Ngāi Tahu, Te Ngāi Tūāhuriri Rūnanga, Te Rūnanga o Arowhenua, Te Rūnanga o Waihao, and Te Rūnanga o Moeraki. This research was funded by a Royal Society of New Zealand Rutherford Discovery Fellowship to MK, and NJF was supported by a University of Otago Doctoral Scholarship. We are grateful to the Aotearoa New Zealand Department of Conservation’s Kakī Recovery Team, Richard Jakob-Hoff at Auckland Zoo, David McLelland at Adelaide Zoo, and John Berry for assisting with provision of kakī, Australian pied stilt, poaka, and interspecific hybrid samples for use in this project, along with associated metadata. We thank our GBS provider AgResearch Ltd, along with thank Olga Kardailsky for assistance in the lab, Dr Alana Alexander for assistance with SLURM job parallelisation, and Jana Wold and two anonymous reviewers for thoughtful feedback on the manuscript.

---

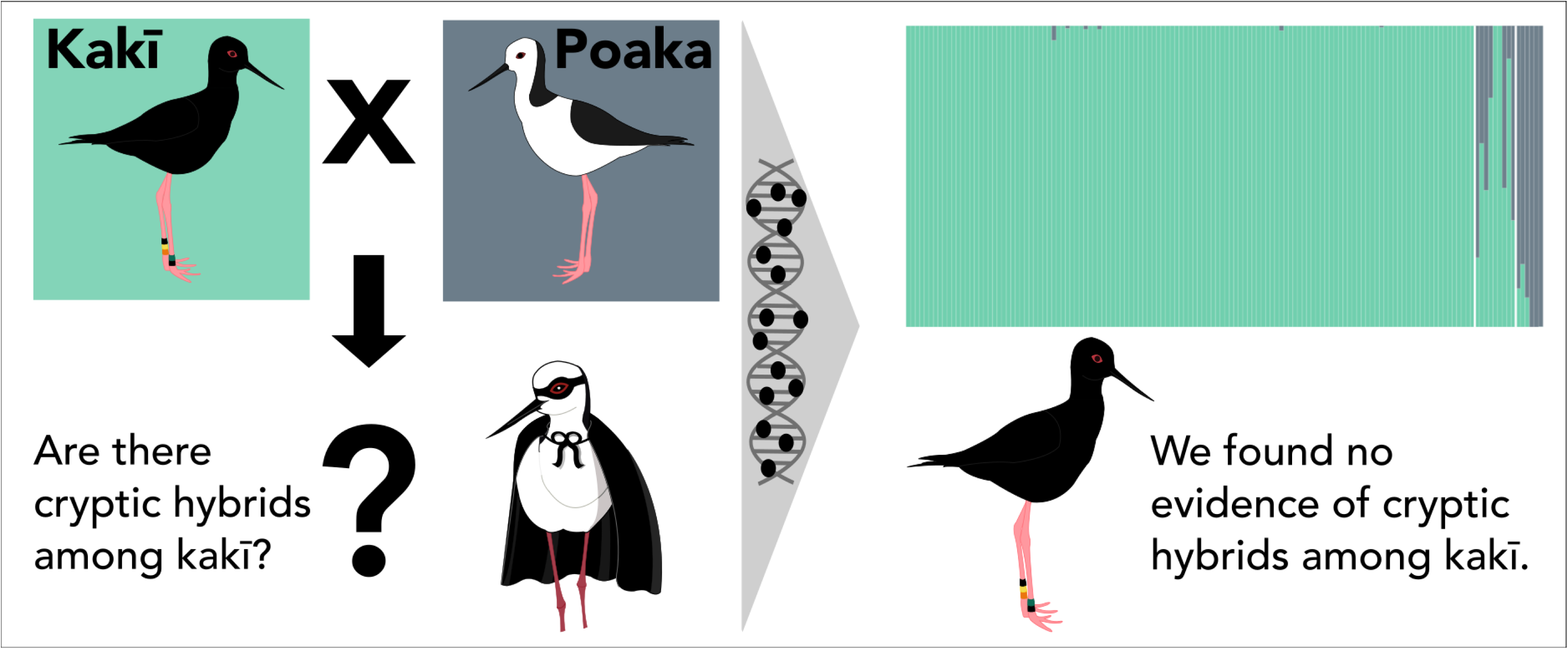

## Notes

### Competing Interest Statement

The authors have declared no competing interest.

### Summary of Updates

Minor change to the title, url corrected in the 'Data Accessibility' section.

