## Supplementary File 1 for "Genomic sequencing confirms absence of introgression despite past hybridisation between a common and a critically endangered bird and its common congener"

### SNPRelate analysis of stilts

Nat Forsdick

18 May, 2021

#### Introduction

This report is supplementary to the manuscript by Forsdick, Martini, Brown, Cross, Maloney, Steeves, Knapp, entitled 'Genomic sequencing confirms absence of introgression despite past hybridisation between a common and a critically endangered bird'. The original Rmarkdown file used to produce this report is available at https://github.com/natforsdick/Himantopus.

##### Session Information

```
## SNPRelate -- supported by Streaming SIMD Extensions 2 (SSE2)
```

```
## R version 3.6.3 (2020-02-29)
## Platform: x86_64-apple-darwin15.6.0 (64-bit)
## Running under: macOS Catalina 10.15.7
##
## Matrix products: default
## BLAS:   /Library/Frameworks/R.framework/Versions/3.6/Resources/lib/libRblas.0.dylib
## LAPACK: /Library/Frameworks/R.framework/Versions/3.6/Resources/lib/libRlapack.dylib
##
## locale:
## [1] en_NZ.UTF-8/en_NZ.UTF-8/en_NZ.UTF-8/C/en_NZ.UTF-8/en_NZ.UTF-8
##
## attached base packages:
## [1] stats     graphics  grDevices utils     datasets  methods   base
##
## other attached packages:
## [1] gridExtra_2.3      pals_1.7           Manu_0.0.1         RColorBrewer_1.1-2
## [5] MASS_7.3-53.1      SNPRelate_1.20.1   gdsfmt_1.22.0
##
## loaded via a namespace (and not attached):
##  [1] digest_0.6.27     grid_3.6.3        gtable_0.3.0      magrittr_2.0.1
##  [5] evaluate_0.14     rlang_0.4.10      stringi_1.5.3     mapproj_1.2.7
##  [9] rmarkdown_2.7     tools_3.6.3       dichromat_2.0-0   stringr_1.4.0
## [13] maps_3.3.0        xfun_0.22         yaml_2.2.1        compiler_3.6.3
## [17] colorspace_2.0-0  htmltools_0.5.1.1 knitr_1.32
```

```
##
## To cite gdsfmt in publications use:
##
##   Xiuwen Zheng, David Levine, Jess Shen, Stephanie M. Gogarten, Cathy
##   Laurie, Bruce S. Weir. A High-performance Computing Toolset for
##   Relatedness and Principal Component Analysis of SNP Data.
##   Bioinformatics 2012; doi: 10.1093/bioinformatics/bts606
```

```
##
##   Xiuwen Zheng, Stephanie M. Gogarten, Michael Lawrence, Adrienne
##   Stilp, Matthew P. Conomos, Bruce S. Weir, Cathy Laurie, David Levine.
##   SeqArray -- A storage-efficient high-performance data format for WGS
##   variant calls. Bioinformatics 2017; doi:
##   10.1093/bioinformatics/btx145
##
## To see these entries in BibTeX format, use 'print(<citation>,
## bibtex=TRUE)', 'toBibtex(.)', or set
## 'options(citation.bibtex.max=999)'.
```

```
##
## To cite SNPRelate in publications use:
##
##   Xiuwen Zheng, David Levine, Jess Shen, Stephanie M. Gogarten, Cathy
##   Laurie, Bruce S. Weir. A High-performance Computing Toolset for
##   Relatedness and Principal Component Analysis of SNP Data.
##   Bioinformatics 2012; doi: 10.1093/bioinformatics/bts606
##
## A BibTeX entry for LaTeX users is
##
##   @Article{,
##     title = {A High-performance Computing Toolset for Relatedness and Principal Component Analysis o
##     author = {Xiuwen Zheng and David Levine and Jess Shen and Stephanie Gogarten and Cathy Laurie an
##     journal = {Bioinformatics},
##     year = {2012},
##     doi = {10.1093/bioinformatics/bts606},
##     volume = {28},
##     number = {24},
##     pages = {3326-3328},
##   }
```

```
##
## To cite the MASS package in publications use:
##
##   Venables, W. N. & Ripley, B. D. (2002) Modern Applied Statistics with
##   S. Fourth Edition. Springer, New York. ISBN 0-387-95457-0
##
## A BibTeX entry for LaTeX users is
##
##   @Book{,
##     title = {Modern Applied Statistics with S},
##     author = {W. N. Venables and B. D. Ripley},
##     publisher = {Springer},
##     edition = {Fourth},
##     address = {New York},
##     year = {2002},
##     note = {ISBN 0-387-95457-0},
##     url = {https://www.stats.ox.ac.uk/pub/MASS4/},
##   }
```

Here we use the SNPRelate package (citation provided in 'Session Information' above) implemented in R version 3.6.3 (2020-02-29) to assess clustering as an alternative approach to DAPC, using the filtered SNP set with all individuals as input ('Forsdick_et_al_Filtered_SNP_set.vcf'). We also generate plots of relatedness within populations, and Fst estimates between populations. The analyses here closely follow the tutorial provided for SNPRelate (https://bioconductor.org/packages/release/bioc/vignettes/SNPRelate/inst/doc/SNPRelate.html).

```
## Principal Component Analysis (PCA) on genotypes:
## Excluding 0 SNP (monomorphic: TRUE, MAF: NaN, missing rate: NaN)
## Working space: 140 samples, 15,851 SNPs
##     using 4 (CPU) cores
## PCA:    the sum of all selected genotypes (0,1,2) = 4035875
## CPU capabilities: Double-Precision SSE2
## Tue May 18 19:36:24 2021    (internal increment: 10968)
## [..................................................]  0%, ETC: ---        [==========================
## Tue May 18 19:36:24 2021    Begin (eigenvalues and eigenvectors)
## Tue May 18 19:36:24 2021    Done.
```

```
## [1] 34.37  5.77  5.25  4.70  4.28  4.08
```

```
##                           sample.id pop         EV1           EV2
## 1 sorted_SQ0683_CCHB8ANXX_s_1_100fq   K  0.02299749 -0.0048721408
## 2 sorted_SQ0683_CCHB8ANXX_s_1_101fq   H -0.28321686  0.1397212112
## 3 sorted_SQ0683_CCHB8ANXX_s_1_102fq   K  0.02612997  0.0009661749
## 4 sorted_SQ0683_CCHB8ANXX_s_1_103fq   K  0.02707698 -0.0035371476
## 5 sorted_SQ0683_CCHB8ANXX_s_1_104fq   K  0.02504148 -0.0024651018
## 6 sorted_SQ0683_CCHB8ANXX_s_1_105fq   K  0.02499737 -0.0021690525
```

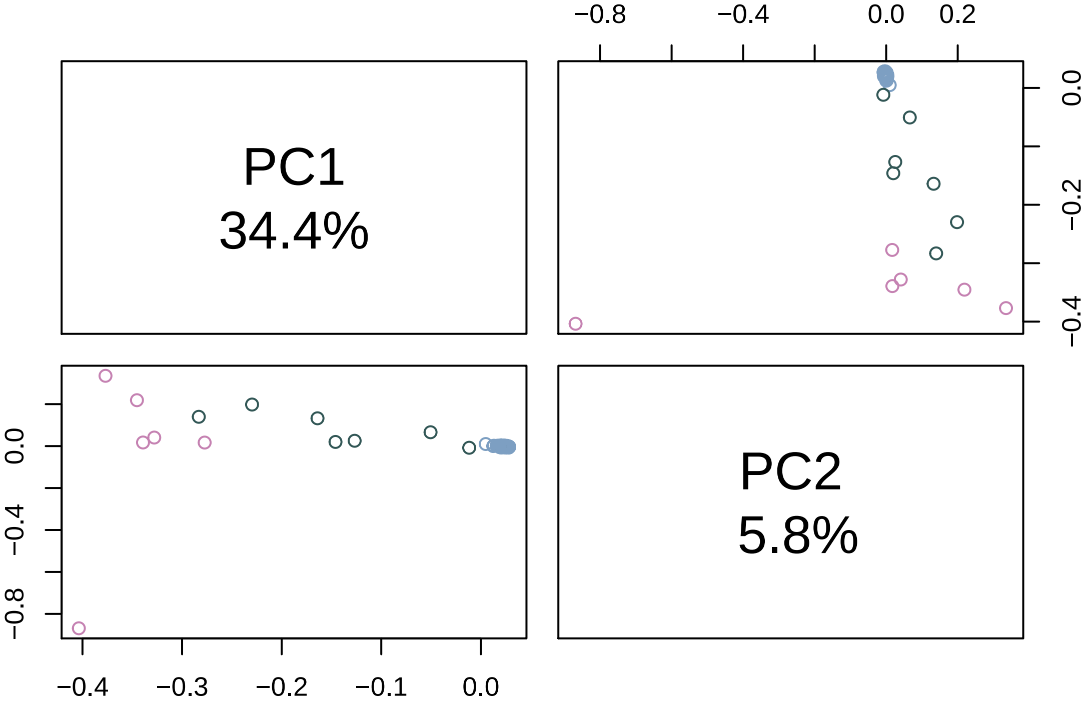


#### Estimating Fst

Here we calculate Fst using the Weir and Cockerham method. First we consider kakī, pied stilts, and hybrids as three distinct units (although they represent two distinct species and intermediate hybrids). The first value represents the weighted Fst estimate, the second is the mean Fst estimate.

```
## Fst estimation on genotypes:
## Excluding 0 SNP (monomorphic: TRUE, MAF: NaN, missing rate: NaN)
## Working space: 140 samples, 15,851 SNPs
## Method: Weir & Cockerham, 1984
```

```
## # of Populations: 3
##     H (9), K (125), P (6)
```

```
## [1] 0.4374895
```

```
## [1] 0.4748272
```

```
##    Min. 1st Qu.  Median    Mean 3rd Qu.    Max.
## -0.0581  0.3130  0.5335  0.4748  0.6853  0.9598
```

Next we only consider kakī and pied stilts. We expect this value to align with our previous estimates produced with VCFTools, as in Table 3 of the main manuscript, where mean Fst = 0.622, and weighted mean Fst = 0.637.

```
## Fst estimation on genotypes:
## Excluding 122 SNPs (monomorphic: TRUE, MAF: NaN, missing rate: NaN)
## Working space: 131 samples, 15,729 SNPs
## Method: Weir & Cockerham, 1984
## # of Populations: 2
##     K (125), P (6)
```

```
## [1] 0.6348234
```

```
## [1] 0.6215057
```

```
##    Min. 1st Qu.  Median    Mean 3rd Qu.    Max.
## -0.3074  0.4707  0.7713  0.6215  0.8478  1.0000
```

#### Relatedness

Now we can look at relatedness metrics. First we are considering Identity-By-Descent, using PLINK method of moment analysis.

First we calculate for kakī:

```
## IBD analysis (PLINK method of moment) on genotypes:
## Excluding 9,122 SNPs (monomorphic: TRUE, MAF: NaN, missing rate: NaN)
## Working space: 125 samples, 6,729 SNPs
##     using 4 (CPU) cores
## PLINK IBD:    the sum of all selected genotypes (0,1,2) = 1471893
## Tue May 18 19:36:25 2021    (internal increment: 65536)
## [..................................................]  0%, ETC: ---        [========================
## Tue May 18 19:36:25 2021    Done.
```

```
##                                  ID1                              ID2        k0
## 1 sorted_SQ0683_CCHB8ANXX_s_1_100fq sorted_SQ0683_CCHB8ANXX_s_1_102fq 1.0000000
## 2 sorted_SQ0683_CCHB8ANXX_s_1_100fq sorted_SQ0683_CCHB8ANXX_s_1_103fq 0.6780768
## 3 sorted_SQ0683_CCHB8ANXX_s_1_100fq sorted_SQ0683_CCHB8ANXX_s_1_104fq 0.8418542
## 4 sorted_SQ0683_CCHB8ANXX_s_1_100fq sorted_SQ0683_CCHB8ANXX_s_1_105fq 1.0000000
## 5 sorted_SQ0683_CCHB8ANXX_s_1_100fq sorted_SQ0683_CCHB8ANXX_s_1_106fq 0.7939919
## 6 sorted_SQ0683_CCHB8ANXX_s_1_100fq sorted_SQ0683_CCHB8ANXX_s_1_107fq 1.0000000
##          k1    kinship
## 1 0.0000000 0.00000000
## 2 0.3219232 0.08048080
## 3 0.1581458 0.03953645
## 4 0.0000000 0.00000000
## 5 0.2060081 0.05150202
## 6 0.0000000 0.00000000
```

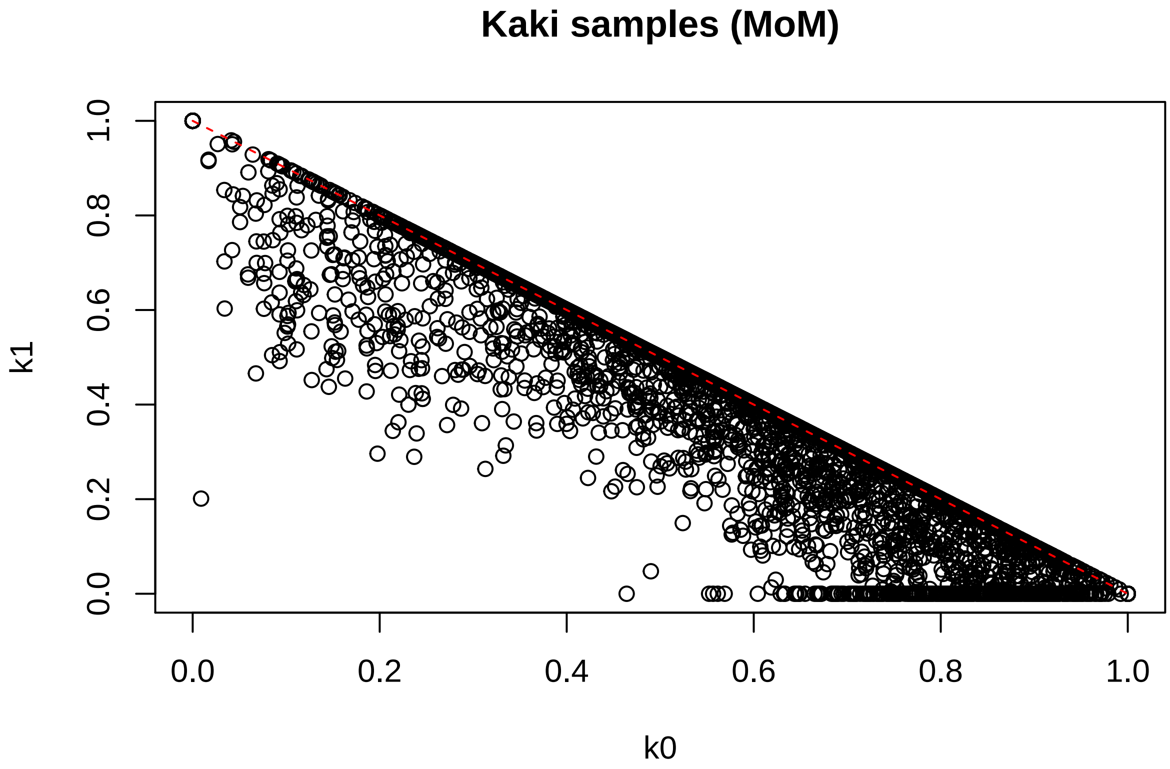


And then for hybrids:

```
## IBD analysis (PLINK method of moment) on genotypes:
## Excluding 2,230 SNPs (monomorphic: TRUE, MAF: NaN, missing rate: NaN)
## Working space: 9 samples, 13,621 SNPs
##     using 4 (CPU) cores
## PLINK IBD:    the sum of all selected genotypes (0,1,2) = 183998
## Tue May 18 19:36:26 2021    (internal increment: 65536)
## [..................................................]  0%, ETC: ---        [==========================
## Tue May 18 19:36:26 2021    Done.
```

```
##                                  ID1                                ID2        k0
## 1 sorted_SQ0683_CCHB8ANXX_s_1_101fq  sorted_SQ0683_CCHB8ANXX_s_1_10fq 1.0000000
## 2 sorted_SQ0683_CCHB8ANXX_s_1_101fq sorted_SQ0683_CCHB8ANXX_s_1_124fq 0.6510915
## 3 sorted_SQ0683_CCHB8ANXX_s_1_101fq sorted_SQ0683_CCHB8ANXX_s_1_136fq 1.0000000
## 4 sorted_SQ0683_CCHB8ANXX_s_1_101fq sorted_SQ0683_CCHB8ANXX_s_1_137fq 1.0000000
## 5 sorted_SQ0683_CCHB8ANXX_s_1_101fq sorted_SQ0683_CCHB8ANXX_s_1_144fq 1.0000000
## 6 sorted_SQ0683_CCHB8ANXX_s_1_101fq sorted_SQ0683_CCHB8ANXX_s_1_149fq 1.0000000
##          k1    kinship
## 1 0.0000000 0.00000000
## 2 0.3489085 0.08722714
## 3 0.0000000 0.00000000
## 4 0.0000000 0.00000000
## 5 0.0000000 0.00000000
## 6 0.0000000 0.00000000
```

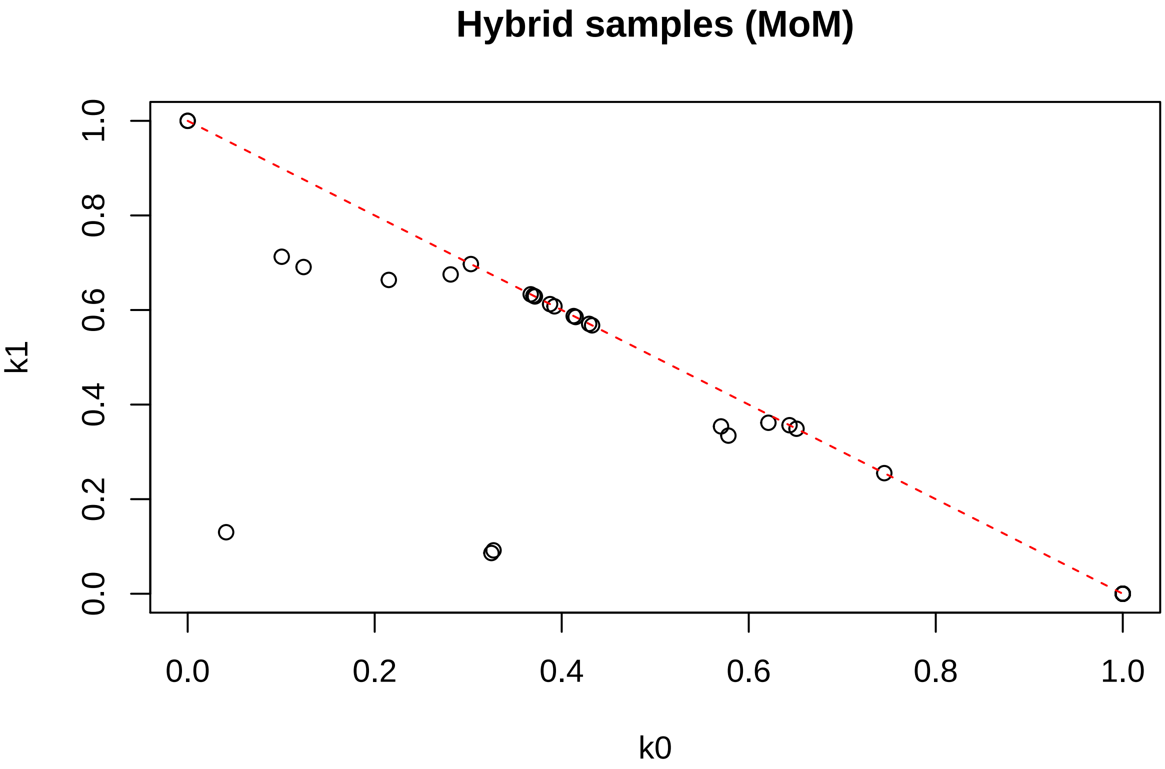


And then for pied stilts:

```
## IBD analysis (PLINK method of moment) on genotypes:
## Excluding 1,152 SNPs (monomorphic: TRUE, MAF: NaN, missing rate: NaN)
## Working space: 6 samples, 14,699 SNPs
##     using 4 (CPU) cores
## PLINK IBD:    the sum of all selected genotypes (0,1,2) = 106652
## Tue May 18 19:36:26 2021    (internal increment: 65536)
## [..................................................]  0%, ETC: ---        [=========================
## Tue May 18 19:36:26 2021    Done.
```

```
##                                  ID1                                ID2        k0
## 1 sorted_SQ0683_CCHB8ANXX_s_1_115fq sorted_SQ0683_CCHB8ANXX_s_1_125fq 0.3756737
## 2 sorted_SQ0683_CCHB8ANXX_s_1_115fq sorted_SQ0683_CCHB8ANXX_s_1_132fq 0.4548257
## 3 sorted_SQ0683_CCHB8ANXX_s_1_115fq sorted_SQ0683_CCHB8ANXX_s_1_139fq 0.5683848
## 4 sorted_SQ0683_CCHB8ANXX_s_1_115fq sorted_SQ0683_CCHB8ANXX_s_1_146fq 0.3851435
## 5 sorted_SQ0683_CCHB8ANXX_s_1_115fq   sorted_SQ0683_CCHB8ANXX_s_1_6fq 0.6880553
## 6 sorted_SQ0683_CCHB8ANXX_s_1_125fq sorted_SQ0683_CCHB8ANXX_s_1_132fq 0.4010705
##          k1    kinship
## 1 0.6243263 0.15608157
## 2 0.5451743 0.13629357
## 3 0.4316152 0.10790380
## 4 0.6148565 0.15371413
## 5 0.3119447 0.07798618
## 6 0.5989295 0.14973236
```

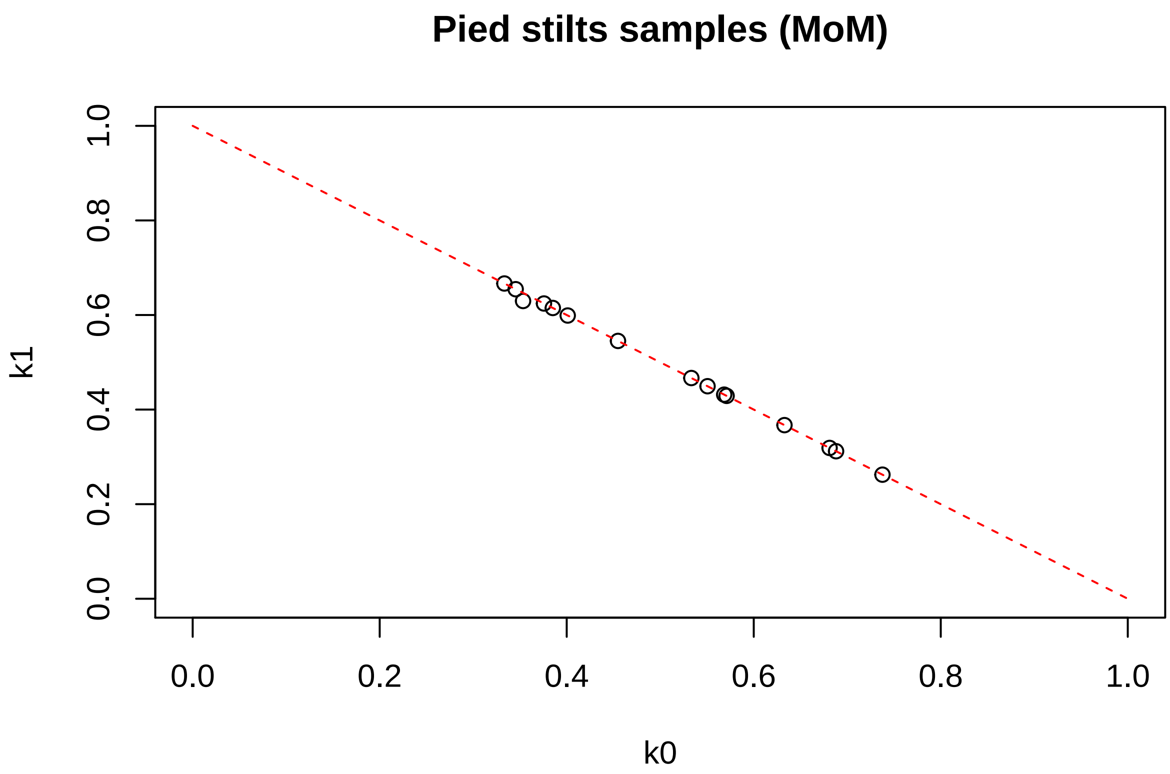


We can also consider all individuals together:

```
## IBD analysis (PLINK method of moment) on genotypes:
## Excluding 0 SNP (monomorphic: TRUE, MAF: NaN, missing rate: NaN)
## Working space: 140 samples, 15,851 SNPs
##     using 4 (CPU) cores
## PLINK IBD:    the sum of all selected genotypes (0,1,2) = 4035875
## Tue May 18 19:36:27 2021    (internal increment: 65536)
## [..................................................]  0%, ETC: ---        [=========================
## Tue May 18 19:36:27 2021    Done.
```

```
##                                ID1                               ID2        k0
## 1 sorted_SQ0683_CCHB8ANXX_s_1_100fq sorted_SQ0683_CCHB8ANXX_s_1_101fq 1.0000000
## 2 sorted_SQ0683_CCHB8ANXX_s_1_100fq sorted_SQ0683_CCHB8ANXX_s_1_102fq 1.0000000
## 3 sorted_SQ0683_CCHB8ANXX_s_1_100fq sorted_SQ0683_CCHB8ANXX_s_1_103fq 0.5189911
## 4 sorted_SQ0683_CCHB8ANXX_s_1_100fq sorted_SQ0683_CCHB8ANXX_s_1_104fq 0.5599956
## 5 sorted_SQ0683_CCHB8ANXX_s_1_100fq sorted_SQ0683_CCHB8ANXX_s_1_105fq 0.5350473
## 6 sorted_SQ0683_CCHB8ANXX_s_1_100fq sorted_SQ0683_CCHB8ANXX_s_1_106fq 0.5634771
##   k1   kinship
## 1  0 0.0000000
## 2  0 0.0000000
## 3  0 0.2405044
## 4  0 0.2200022
## 5  0 0.2324763
## 6  0 0.2182615
```

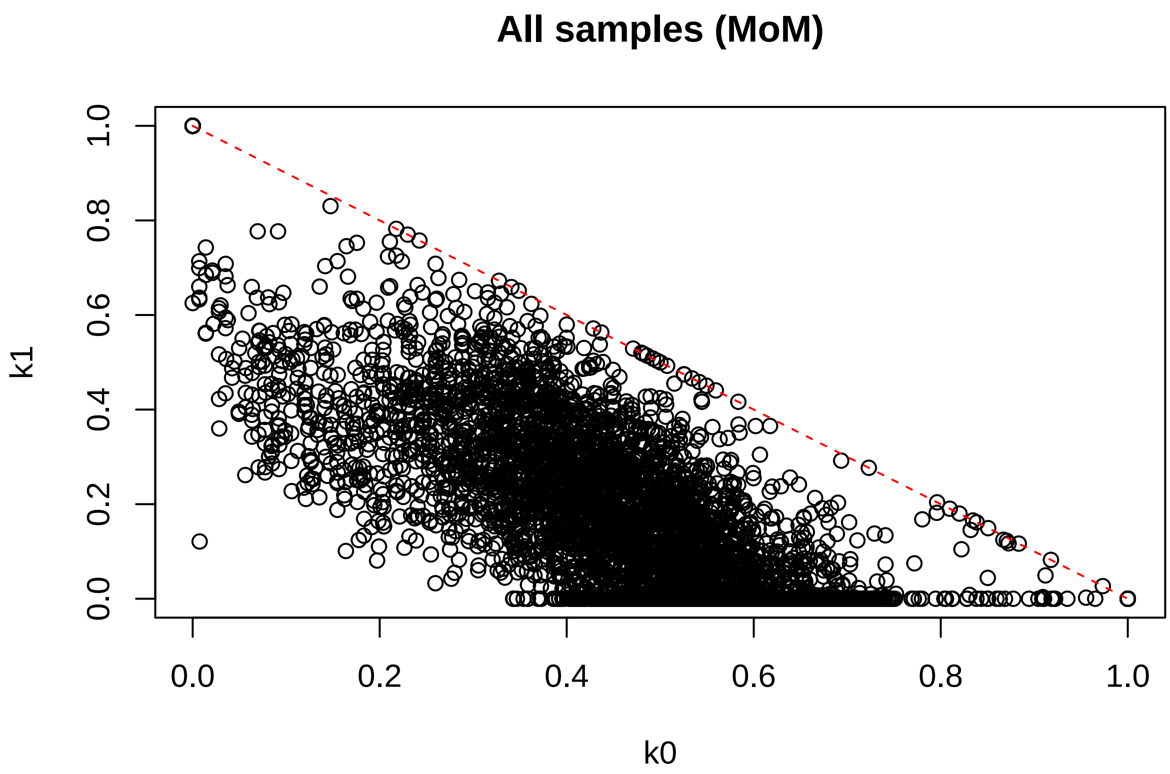


Then we can look at identity-by-state:

```
## Identity-By-State (IBS) analysis on genotypes:
## Excluding 0 SNP (monomorphic: TRUE, MAF: NaN, missing rate: NaN)
## Working space: 140 samples, 15,851 SNPs
##     using 2 (CPU) cores
## IBS:    the sum of all selected genotypes (0,1,2) = 4035875
## Tue May 18 19:36:28 2021    (internal increment: 65536)
## [..................................................]  0%, ETC: ---        [==========================
## Tue May 18 19:36:28 2021    Done.
```

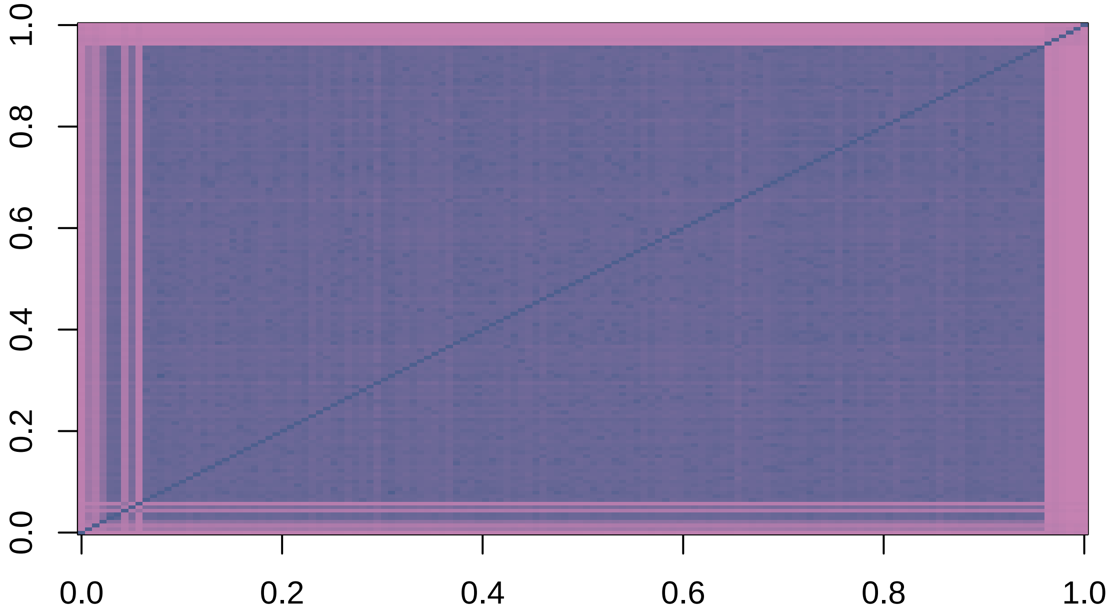

This is more difficult to interpret because individuals are not perfectly clustered by population in the input files.

##### Additional clustering analysis

We can also perform multidimensional scaling analysis on the n × n matrix of genome-wide IBS pairwise distances:

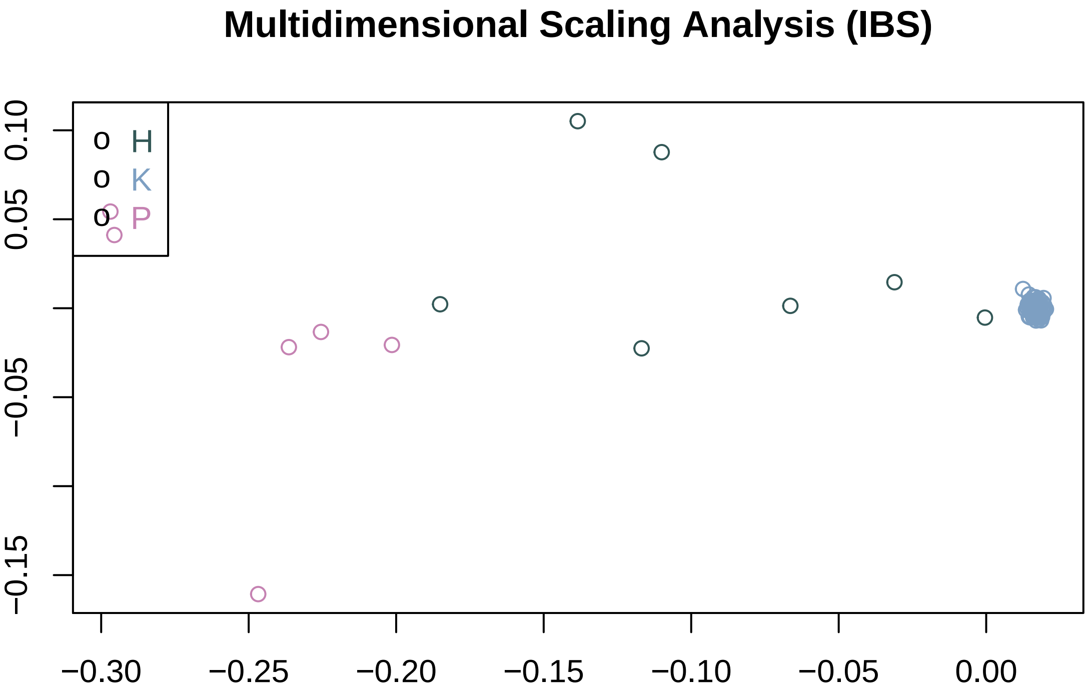


To perform cluster analysis on the n × n matrix of genome-wide IBS pairwise distances, and determine the groups by a permutation score:

```
## Identity-By-State (IBS) analysis on genotypes:
## Excluding 0 SNP (monomorphic: TRUE, MAF: NaN, missing rate: NaN)
## Working space: 140 samples, 15,851 SNPs
##     using 2 (CPU) cores
## IBS:    the sum of all selected genotypes (0,1,2) = 4035875
## Tue May 18 19:36:28 2021    (internal increment: 65536)
## [..................................................]  0%, ETC: ---        [==========================
## Tue May 18 19:36:29 2021    Done.

## Determine groups by permutation (Z threshold: 15, outlier threshold: 5):
## Create 1 groups.

##
## G001
##  140

## Create 3 groups.

##
##   H   K   P
##   9 125   6
```

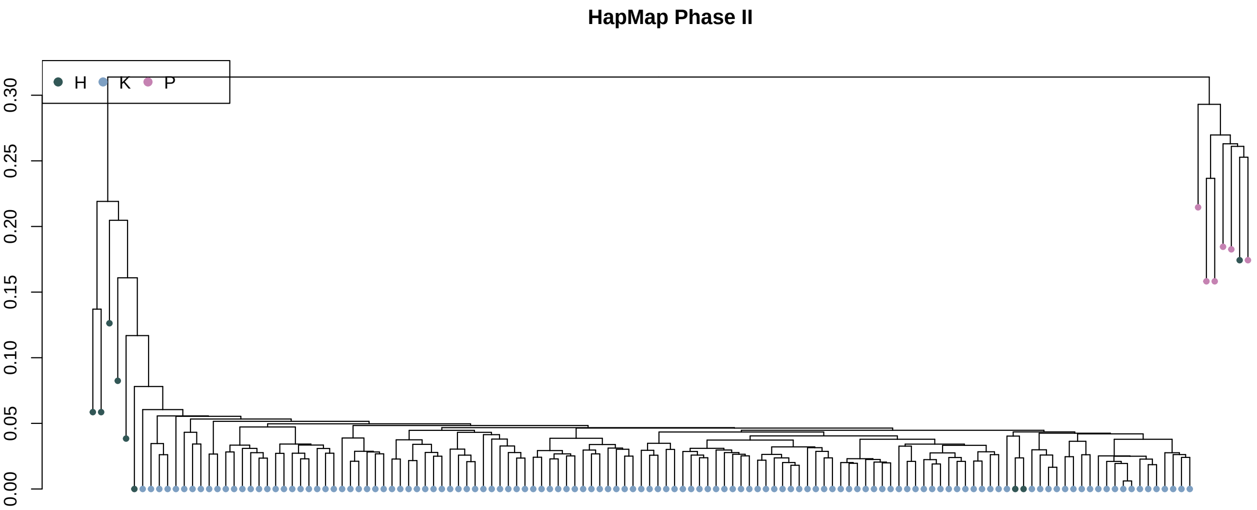


### Conclusions

The PCA results appear concordant to those of DAPC, where kakī and pied stilts are grouped distinctly from one another, with hybrids falling intermediate to the two. The Fst estimates indicate strong differentiation between kakī and pied stilts. When hybrids are included, the differentiation is still strong but less so, as we would expect.

The clustering analysis represented in the 'HapMap Phase II' plot again shows kakī and pied stilts to cluster distinctly, with a handful of hybrids grouping with each of these two groups. Overall, we interpret these results as concordant with thos of DAPC and ADMIXTURE analyses, providing further support for our conclusion that despite a history of hybridisation, there has been no substantial admixture of poaka genetic material into the kakī genome.
